# The identification of mecciRNAs and their roles in mitochondrial entry of proteins

**DOI:** 10.1101/668665

**Authors:** Xu Liu, Xiaolin Wang, Jingxin Li, Shanshan Hu, Yuqi Deng, Hao Yin, Xichen Bao, Qiangfeng Cliff Zhang, Geng Wang, Baolong Wang, Qinghua Shi, Ge Shan

## Abstract

Mammalian mitochondria have small genomes encoding very limited numbers of proteins. Over one thousand proteins and noncoding RNAs encoded by nuclear genome have to be imported from the cytosol into the mitochondria. Here we report the identification of hundreds of circular RNAs (mecciRNAs) encoded by mitochondrial genome. We provide both *in vitro* and *in vivo* evidence to show that mecciRNAs facilitate mitochondrial entry of nuclear-encoded proteins by serving as molecular chaperones in the folding of imported proteins. Known components of mitochondrial protein and RNA importation such as TOM40 and PNPASE interact with mecciRNAs and regulate protein entry. Expression of mecciRNAs is regulated, and these transcripts are critical for mitochondria in adapting to physiological conditions and diseases such as stresses and cancers by modulating mitochondrial protein importation. mecciRNAs and their associated physiological roles add categories and functions to eukaryotic circular RNAs, and shed novel lights on communication between mitochondria and nucleus.

## Introduction

Mammalian mitochondria have their own genome to encode 13 mitochondrial specific proteins and multiple noncoding RNAs such as 16S rRNA and some tRNAs (Anderson et al., 1981; Gustafsson et al., 2016). These proteins and RNAs are a very small fraction of mitochondrial contents, and more than one thousand proteins and an array of noncoding RNAs encoded by the nuclear genome are known to be transported into mammalian mitochondria to sustain the homeostasis and duplication of these cell organelles (Harbauer et al., 2014).

Over one thousand mitochondrial proteins encoded by the nuclear genome are imported into mitochondria through the outer membrane TOM40 complex, with the help of cytosolic chaperones of the heat-shock family proteins (Deshaies et al., 1988; Young et al., 2003). Some nuclear-encoded noncoding RNAs such as MRP RNA (RMRP), 5S rRNA, and certain tRNAs are also imported into mitochondria, although the mitochondrial RNA importation mechanism is much less understood (Smirnov et al., 2010; Wang et al., 2010). It has been known that a polynucleotide phosphorylase (PNPASE) PNPT1 residing in the intermembrane space plays an unanticipated role in mitochondrial RNA importation (Cheng et al., 2018; Wang et al., 2010).

Essentially all mitochondrial functions rely on import of proteins and RNAs, and cells have to manage the importation according to different physiological conditions (Harbauer et al., 2014; Quirós et al., 2015; Smirnov et al., 2010; Sokol et al., 2014). When mitochondrial functions are compromised, nuclear responses such as mtUPR (mitochondrial unfolded protein response), UPRam (unfolded protein response activated by mistargeting of proteins), and mPOS (mitochondrial precursor over-accumulation stress) are triggered to express mitochondrial chaperones, to induce the degradation of unimported proteins, and to reduce mRNA translation in the cytosol (Boos et al., 2019; Scheibye-Knudsen et al., 2015; Topf et al., 2016; Wang and Chen, 2015; Wrobel et al., 2015). Whether and how mitochondria themselves cope with dynamic cellular conditions by regulating protein importation remain much less studied (Moye-Rowley, 2003; Weidberg and Amon, 2018).

In an effort to reveal novel RNAs in mitochondria, we have identified hundreds of circular RNAs (circRNAs) encoded by the mitochondrial genome from both human and murine cells. Recent years have witnessed the identification of thousands of circRNAs encoded by the nuclear genome, and some of them function as ceRNAs (microRNA sponges), protein binding partners, and transcriptional regulators (Chen et al., 2015; Li et al., 2018). Here, we provided lines of evidence to show that some of the mitochondria encoded circRNAs could facilitate the mitochondrial entry of proteins encoded by the nuclear genome. Mitochondria modulated the amount of specific proteins imported into these organelles through mecciRNAs to adjust to physiological conditions.

## RESULTS

### Identification of mitochondria encoded circRNAs

We sequenced RNAs from isolated mitochondria and analyzed the RNA-sequencing (RNA-seq) data with pipelines to identify circRNAs encoded by either the nuclear genome or the mitochondrial genome (Memczak et al., 2013) (Figure 1A). From four cell lines and tissues, 209 and 274 <u>m</u>itochondria-<u>e</u>n<u>c</u>oded <u>ci</u>rcRNAs (mecciRNAs) were identified in human and mice, respectively (Figure 1B and Figure S1A). To expand the search, we also sequenced mitochondrial RNAs from adult male zebrafish (*Danio rerio*), and 139 mecciRNAs were identified (Figure 1B and Figure S1A). Thus, it seemed that mecciRNAs were present in vertebrates. Bioinformatics analysis revealed that some circRNAs encoded by the nuclear genome (g-circRNAs) were also present in the isolated mitochondria, although with significantly lower concentration (RPM) than mecciRNAs (Figure 1B). mecciRNAs and g-circRNAs shared similar AG/GT junction motif (Figure 1C and Figure S1B). No linear splicing event was identified in mitochondrial transcripts. Other features of mecciRNAs such as the 5’ and 3’ flanking sites and the size distribution of mecciRNAs were also analyzed (Figure S1C and S1D).

**Figure 1.**
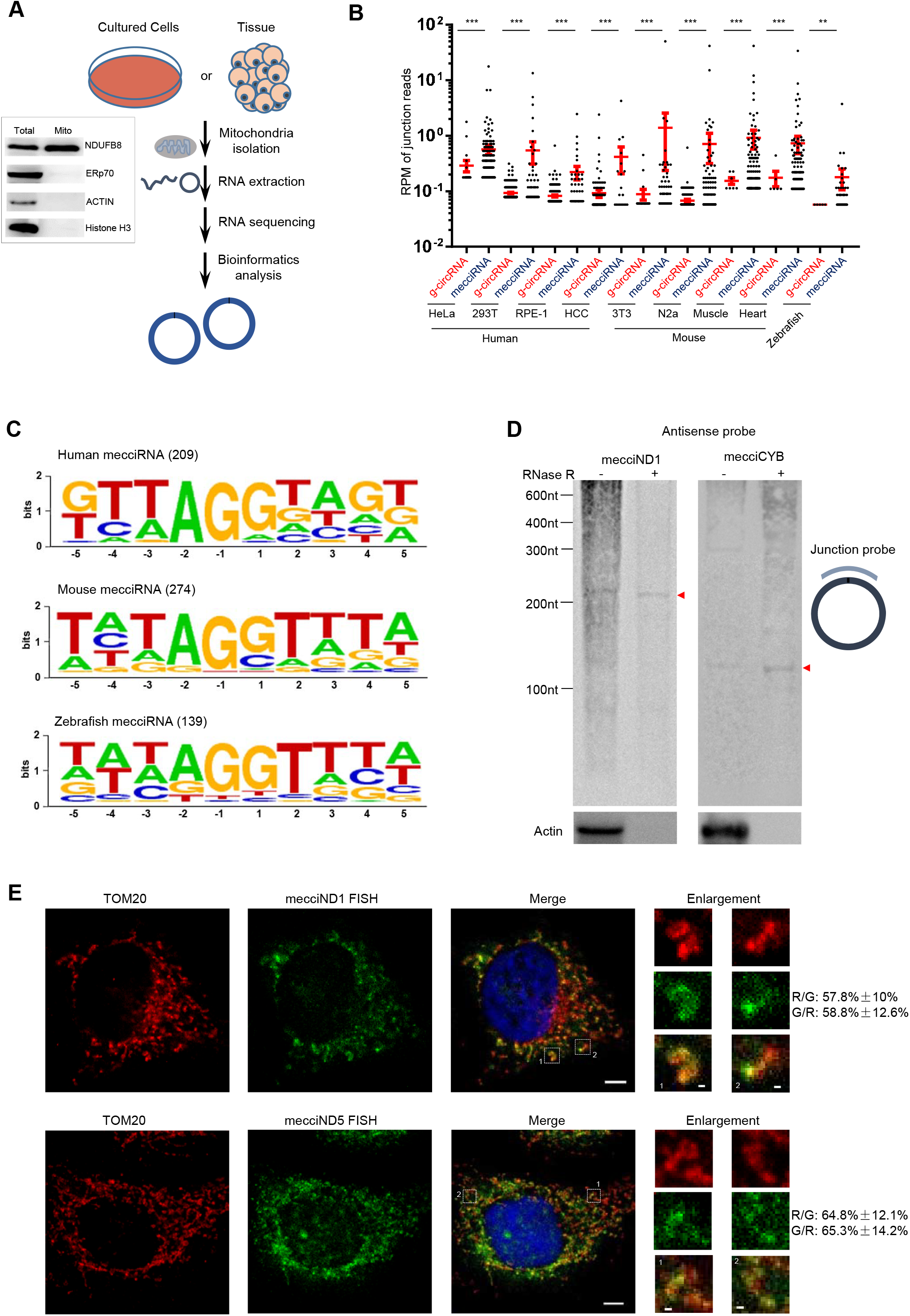
Identification and characterization of mecciRNAs. (A) Experimental procedures for mitochondrial RNA sequencing to identify mecciRNAs; Western blots showing the quality of purified mitochondria are presented; ERp70, ER marker; Histone H3, nuclear marker; ACTIN, cytosolic marker; NDUFB8, a mitochondrial inner membrane protein. (B) g-circRNAs and mecciRNAs (reads ≥ 2) from sequencing data of mitochondrial (mito) RNAs in human and murine cells and tissues; Hepatocellular carcinoma (HCC). (C) Junction motif of human, mouse and zebrafish (adult male) mecciRNAs. (D) Northern blots of mecciRNAs. RNase R treated (+) and untreated (-) total RNAs from HeLa cells were examined; the position of the probe is indicated; actin mRNA served as linear control. (E) Confocal images in single z-section of immunofluorescence (IF) for TOM20 together with FISH of mecciND1 or mecciND5. Boxed areas are enlarged. Colocalization between mecciND1 or mecciND5 (G, green), TOM20 (R, red) is shown (n=20 randomly selected areas). R/G, The proportion of red signal to green signal colocalization; G/R, The proportion of green signal to red signal colocalization. In (E), scale bars, 5 µm and 500 nm (enlarged areas); in (B), error bars, s.e.m.; **P < 0.01; ***P < 0.001 by two-tailed Mann–Whitney U test.

We then verified some of these mecciRNAs with Northern blots and PCR (Figure 1D and Figure S2A-S2E). To exclude potential misidentification of junction sites due to differences in annotated genome and genome of cultured cells, nuclear DNA and mitochondrial DNA from HeLa cells were re-sequenced, and no DNA read matching junction sequences of human mecciRNAs was identified (Figure S3A). In previously published RNA-seq data of cells without mitochondria (rho0 MEF cells), no mecciRNA was identified, whereas mecciRNAs were found in wildtype MEF cells (Shimada et al., 2018) (Figure S3B). Interestingly, considering the small size of mitochondrial genome, mecciRNAs were relatively enriched in nascent circRNAs detected from HeLa cells (Bao et al., 2018) (Figure S3C). Also, thousands of g-circRNAs, but no mecciRNAs were identified in RNA-seq of nuclear RNA samples from isolated nuclei of human HeLa and HEK293T cells as well as murine N2a and 3T3 cells.

Furthermore, mecciRNAs only accounted for a very small portion of all circRNAs in the cytoplasm as g-circRNAs were much more predominant in molecular numbers than mecciRNAs (Figure S4A). In contrast to mitochondria-encoded mRNAs, mecciRNAs were also present outside of the mitochondria (Figure S4B). We then examined two mecciRNAs encoded by the mitochondrial gene ND1 (mecciND1) and ND5 (mecciND5) in details. RNA fluorescence *in situ* hybridization (FISH) combined with immunofluorescence (IF) of TOM20 (mitochondrial outer membrane protein) revealed that both FISH signals of mecciND1 and mecciND5 were partially overlapped with TOM20 IF signals (Figure 1E). A portion of mecciND1 and mecciND5 signals did not overlap with the mitochondria, indicating that some of these mecciRNAs were also present outside of the mitochondria and in the cytosol (Figure 1E). As a comparison, FISH signals of the ND1 mRNAs were largely colocalized with TOM20 IF signals (Figure S4C).

### Interaction between mecciND1 and RPA proteins

We decided to investigate potential functions of mecciRNAs using mecciND1. Pulldown of mecciND1 with antisense probe was performed, and two proteins, RPA70 and RPA32 were identified to be co-pulled down with this circRNA (Figure 2A and Figure S5A). The interaction between mecciND1 and RPA70 & RPA32 proteins was then confirmed with RNA pulldown followed by Western blotting and protein immunoprecipitation followed by RNA detection (RNA-IP, or RIP) (Figure 2B-2D and Figure S5B). FISH of mecciND1 along with IF of RPA proteins and TOM20 showed that over half of the mecciND1 signals overlapped with the mitochondria and cytoplasmic RPA proteins (Figure S5C).

**Figure 2.**
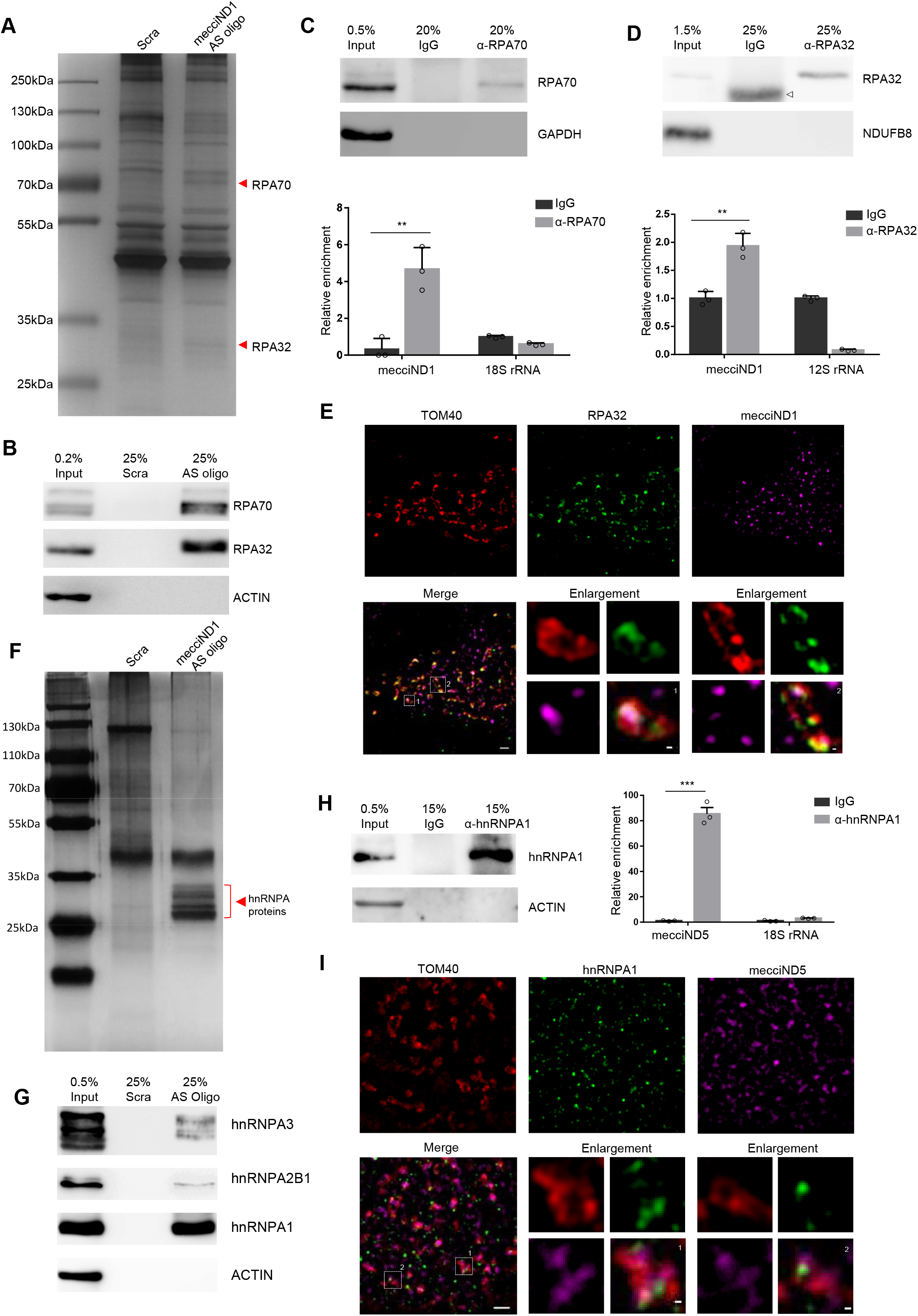
mecciND1 interacts with RPA70 & RPA32 and mecciND5 interacts with hnRNPA proteins. (A) Pulldown of MecciND1 with biotin labeled antisense oligo (AS oligo) in HeLa cells. Proteins co-pulled down with mecciND1 were subjected to silver staining; red triangles indicate the bands identified as RPA70 and RPA32 by mass spectrometry. Scra, oligos with scrambled sequences. (B) RPA70 and RPA32 co-pulled down with mecciND1 were verified by Western blots; ACTIN, negative control. (C) RNA immunoprecipitation (RIP) against RPA70 (α-RPA70) with whole cell lysate of HeLa cells. Successful IP of the protein was detected by Western blots; GAPDH, negative control. RNAs from RIP were quantified by real-time qPCR; 18S rRNA, negative control. (D) RNA immunoprecipitation (RIP) against RPA32 (α-RPA32) with mitochondrial lysate of HeLa cells. Successful IP of the protein was detected by Western blots; NDUFB8, a mitochondrial marker and the negative control for IP; there was a non-specific band (light chain) in the IgG control (indicated with an open triangle). RNAs from RIP were quantified by real-time qPCR; mitochondrial 12S rRNA, negative control. (E) Representative structured illumination microscopy images (N-SIM) in single z-section of immunofluorescence (IF) for PRA32 together with TOM40 as well as FISH of mecciND1 in fixed HeLa cells. Boxed areas are enlarged; scale bars, 2 µm and 200 nm (enlarged areas). (F) Pulldown of mecciND5 with biotin labeled antisense oligo in HeLa cells. Proteins co-pulled down with mecciND5 were subjected to silver staining; red triangles indicate the bands identified as hnRNPA1, hnRNPA2B1 and hnRNPA3 by mass spectrometry. Scra, oligos with scrambled sequences. (G) hnRNPA1, hnRNPA2B1, and hnRNPA3 co-pulled down with mecciND5 were verified by Western blots; ACTIN, negative control. (H) RNA immunoprecipitation (RIP) against hnRNPA1 (α-hnRNPA1) with whole cell lysate of HeLa cells. Successful IP of the protein was detected by Western blots; ACTIN, negative control. RNAs from RIP were quantified by real-time qPCR; Actin mRNA, negative control. (I) Representative structured illumination microscopy (N-SIM) images in single z-section of immunofluorescence (IF) for hnRNPA1 together with TOM40 as well as FISH of mecciND5 in fixed HeLa cells. Scale bars, 2 µm and 200 nm (Boxed areas are enlarged). In (C), (D) and (H) error bars, s.e.m.; n=3 independent experiments; **P < 0.01, ***P < 0.001, Student’s *t*-test.

We next examined the localization of TOM40, RPA32, and mecciND1 in greater detail with super-resolution N-SIM (Figure 2E and Figure S5D). At the resolution of SIM, we could differentiate whether fluorescent signals were from inside or outside of the mitochondrion (the TOM40 enclosed space). We observed that RPA32 and mecciND1 had localizations inside of the TOM40 enclosed space, and RPA32 and mecciND1 had colocalizations around or inside TOM40 signals (Figure 2E and Figure S5D). A fraction of RPA32 and mecciND1 also had localizations outside of the mitochondria (Figure 2E and Figure S5D).

### Interaction between mecciND5 and hnRNPA proteins

We next investigated whether another mecciRNA, mecciND5, could interact with specific proteins. Antisense probe pulldown of mecciND5 revealed three proteins: hnRNPA1, hnRNPA2B1, and hnRNPA3 to be co-pulled down (Figure 2F and Figure S6A). RNA pulldown of mecciND5 followed by Western blotting detected hnRNPA1, hnRNPA2B1, and hnRNPA3 (collectively hnRNPA proteins) (Figure 2G). hnRNPA1, hnRNPA2B1, and hnRNPA3 proteins share high homology, and hnRNPA1 was also the predominant one among hnRNPA proteins interacting with mecciND5 (Figure 2F, 2G and Figure S6B). RNA-IP with antibodies against hnRNPA1 showed enrichment of mecciND5 (Figure 2H). IF of hnRNPA proteins and TOM20 revealed that more than half of the cytoplasmic hnRNPA proteins colocalized with TOM20 (Figure S6C). More than half of cytoplasmic hnRNPA IF signals also colocalized with cytoplasmic mecciND5 signals (Figure S6C). With super-resolution N-SIM, we found that hnRNPA1 and mecciND5 had localizations inside of the TOM40 enclosed space, and hnRNPA1 and mecciND5 had colocalizations around TOM40 signals (Figure 2I and Figure S6D). Some hnRNPA1 and mecciND5 also localized outside of the mitochondria (Figure 2I and Figure S6D).

### Mitochondrial RPA levels were regulated by mecciND1

Interestingly, knockdown of mecciND1 with siRNAs decreased RPA70 and RPA32 protein levels in mitochondria, albeit the total levels of both proteins were less affected (Figure 3A and Figure S7A). In another attempt to interfere with mecciND1, we applied antisense morpholino oligos (AMO) against mecciND1 (Figure S7B). Surprisingly, mitochondrial RPA protein (especially RPA32) levels were increased after mecciND1-AMO transfection (Figure S7B). We found that whole cell levels of mecciND1 were not changed, but cytosolic mecciND1 levels were increased (Figure S7B). This result suggested that mecciND1-AMO somehow held mecciND1 in the cytosol, and the higher cytosolic mecciND1 levels resulted in more RPA32 importation into mitochondria. We next found that mecciND1 could be overexpressed with a nuclear transfected plasmid (Figure S7C). Overexpression of mecciND1 with nuclear plasmids increased RPA70 and RPA32 protein levels in mitochondria, and again the total levels of both proteins were not significantly changed (Figure 3B). Either mecciND1 knockdown or overexpression did not affect mRNA levels of RPA70 and RPA32 (Figure S7D). When examined with N-SIM, RPA32 signals inside TOM40 enclosed space were significantly decreased in the mecciND1 knockdown cells as compared to those of control cells (Figure 3C, 3D and Figure S7E, S7F).

**Figure 3.**
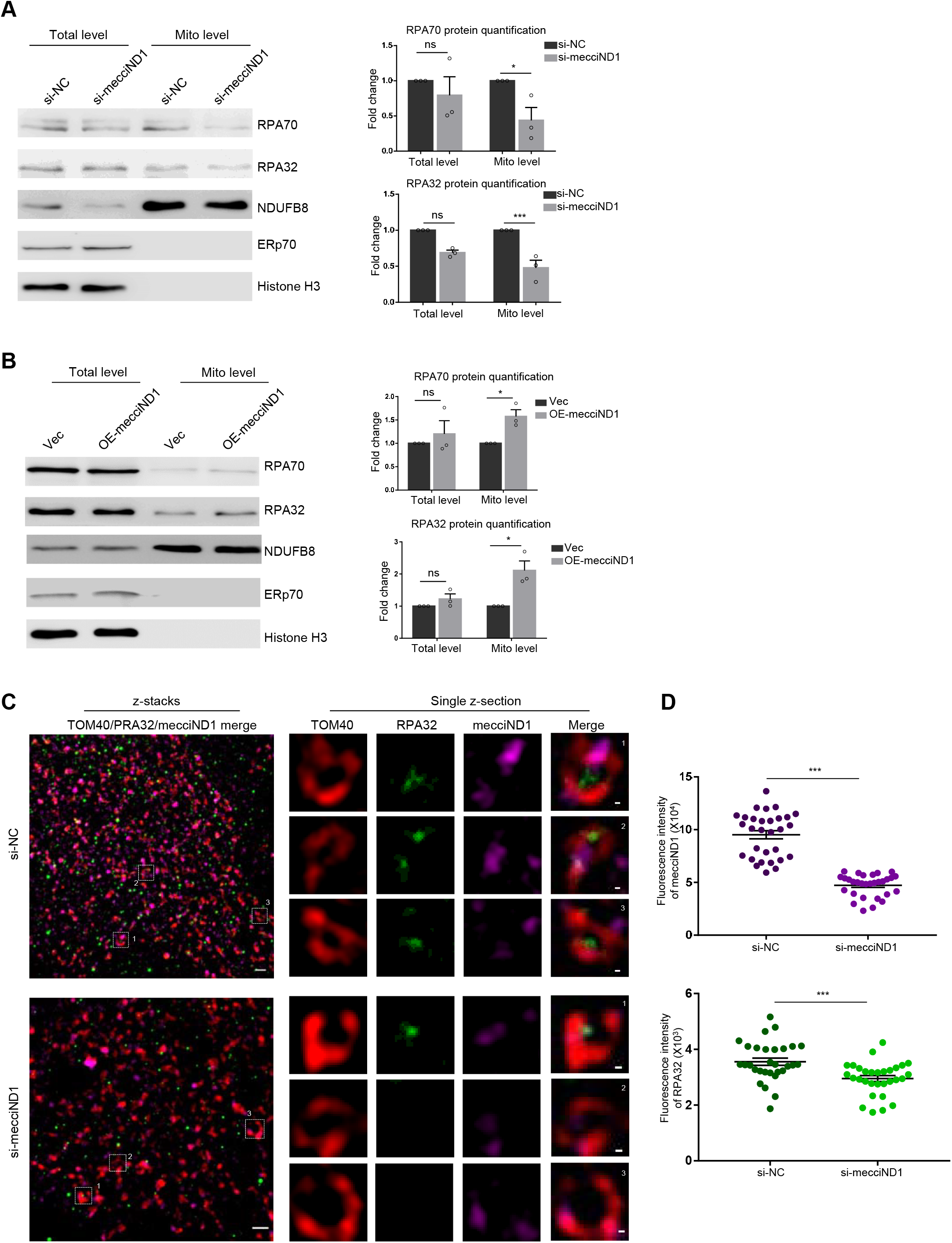
mecciND1 regulates mitochondrial RPA levels. (A) Mitochondrial RPA70 and RPA32 protein levels were decreased after knockdown of mecciND1 with siRNA in 293T cells. Western blots of proteins from whole cells (total level) and mitochondria (mito level) were examined; si-NC, siRNA with scrambled sequences; NDUFB8 served as a loading control for mitochondrial proteins; ERp70, an endoplasmic reticulum marker; Histone H3, a nuclear marker. Quantification of RPA proteins is shown (normalized to NDUFB8). (B) Mitochondrial RPA70 and RPA32 protein levels were increased under mecciND1 overexpression in 293T cells. (C) Representative structured illumination microscopy images in z-stacks (3D N-SIM) of immunofluorescence (IF) for PRA32 together with TOM40 as well as FISH of mecciND1 in fixed RPE-1 cells transfected with siRNA (si-NC or si-mecciND1). Images of single z-section of boxed areas are enlarged. Scale bars, 2 µm and 200 nm (enlarged areas). (D) Quantification of fluorescence signals of mecciND1 and RPA32 from N-SIM is shown; areas representing single mitochondrion were selected and fluorescence signals overlapping the TOM40 signal or inside the TOM40 enclosed space were quantified; n=30 randomly chosen areas. In (A), (B) and (D), error bars, s.e.m.; in (A) and (B), n=3 independent experiments; ns, not significant; *P < 0.05, **P < 0.01, ***P < 0.001 by Student’s *t*-test.

### Mitochondrial hnRNPA levels were regulated by mecciND5

Knockdown of mecciND5 with siRNAs decreased hnRNPA protein levels in mitochondria, but the total levels of all three hnRNPA proteins and mRNAs were much less affected (Figure 4A and Figure S8A). Mitochondrial hnRNPA protein levels were found to be increased after mecciND5-AMO transfection (Figure S8B). The whole cell levels of mecciND5 were unchanged, but cytosolic mecciND5 levels increased (Figure S8B). Overexpression of mecciND5 with a nuclear expressed plasmid increased hnRNPA1 and hnRNPA2B1 protein levels in mitochondria, and whole cell levels of all hnRNPA proteins and mRNAs were unchanged (Figure 4B and Figure S8C, S8D). When examined with N-SIM, hnRNPA1 signals inside TOM40 enclosed space were significantly decreased in the mecciND5 knockdown cells as compared to those of control cells (Figure 4C, 4D and Figure S8E, S8F).

**Figure 4.**
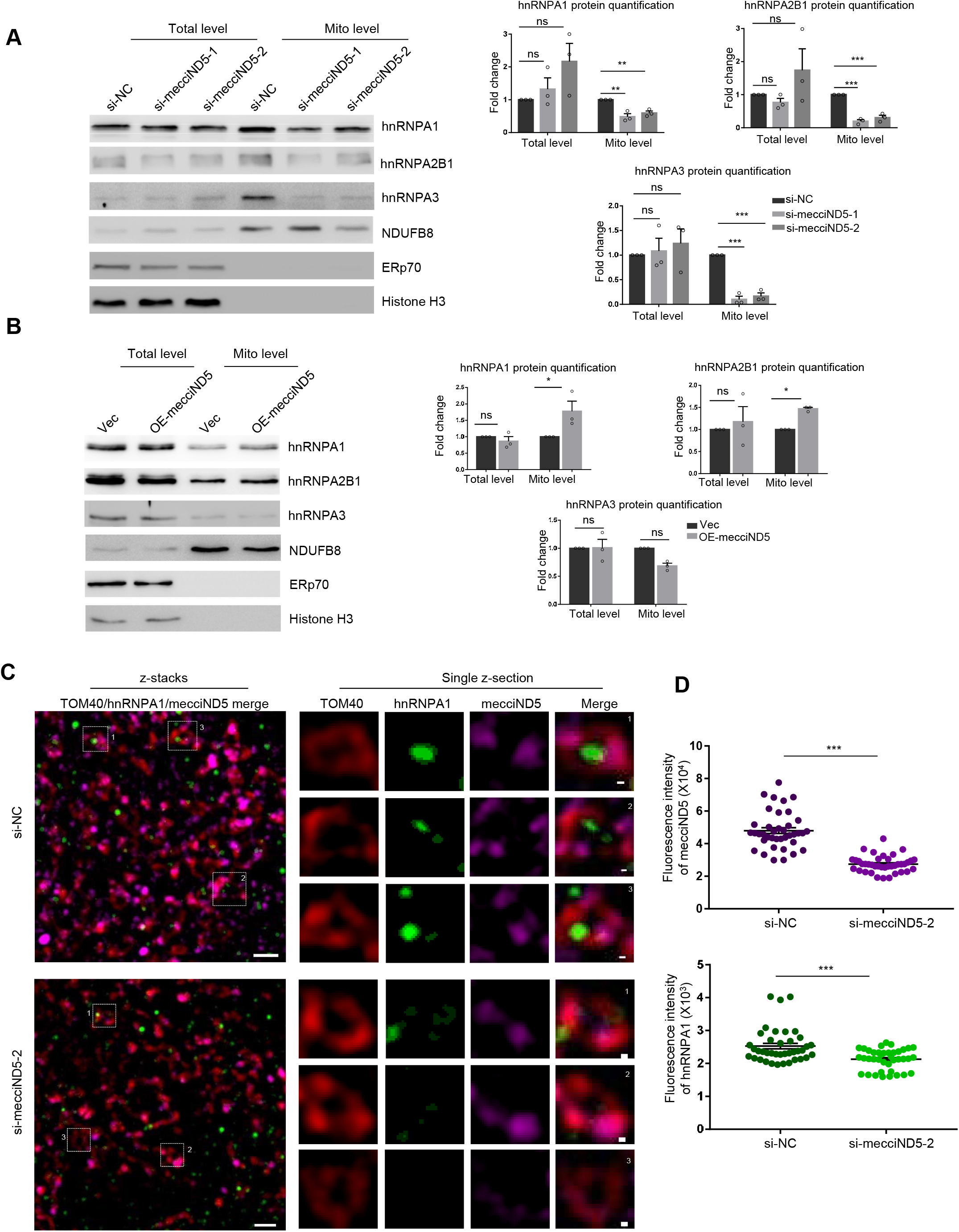
mecciND5 regulates mitochondrial hnRNPA levels. (A) Mitochondrial hnRNPA1, hnRNPA2B1, and hnRNPA3 protein levels were decreased after knockdown of mecciND5 with siRNAs in 293T cells. Western blots of proteins from whole cells (total level) and mitochondria (mito level) are shown; si-NC, siRNA with scrambled sequences; NDUFB8 served as a loading control for mitochondrial proteins; ERp70, an endoplasmic reticulum marker; Histone H3, a nuclear marker. Quantification of hnRNPA proteins is shown (normalized to NDUFB8). (B) Mitochondrial hnRNPA1, hnRNPA2B1, and hnRNPA3 protein levels were increased under mecciND5 overexpression in 293T cells. (C) Representative structured illumination microscopy images in z-stacks (3D N-SIM) of immunofluorescence (IF) for hnRNPA1 together with TOM40 as well as FISH of mecciND5 in fixed RPE-1 cells transfected with siRNA (si-NC or si-mecciND5). Single z-section image of boxed areas is enlarged. Scale bars, 2 µm and 200 nm (enlarged areas). (D) Quantification of fluorescence signals of mecciND5 and hnRNPA1 from N-SIM is shown; areas representing single mitochondrion were selected and fluorescence signals overlapped with the TOM40 signal or inside the TOM40 enclosed space were quantified; n=39 randomly chosen areas. In (A), (B) and (D), error bars, s.e.m; in (A) and (B), n=3 independent experiments; ns, not significant; *P < 0.05; **P < 0.01; ***P < 0.001, Student’s *t*-test.

Taken together, for both mecciND1 and mecciND5, their cellular levels were positively correlated with levels of their corresponding protein partners inside mitochondria, and these data suggested a possibility that both mecciRNAs promoted mitochondria importation of specific proteins. The abundance of these RNAs was reasonable for them to play this role, as there were ~52-109 molecules of mecciND1 and ~118-330 molecules of mecciND5 in each cell in the four cell lines we examined (Figure S9A).

### mecciRNAs promote mitochondrial importation in *in vitro* assays

We then performed a series of assays with purified mitochondria and *in vitro* transcribed linear and circular RNA (Wesselhoeft et al., 2018) (Figure S9B, S9C). Through RNA import assay, we found that extra-mitochondrial mecciND1 and mecciND5 but not the g-circRNA circSRSF could be imported into mitochondria, and all their linear forms could not get into mitochondria (Figure 5A). When mecciND1 added together with RPA32 mRNA in the rabbit reticulocyte translational system (to examine co-translational effect of mecciRNA), mecciND1 increased the mitochondrial importation of *in vitro* translated RPA32 protein (Figure 5B and Figure S9D). Similarly, co-translationally added mecciND5 also increased the mitochondrial importation of *in vitro* translated hnRNPA1 (Figure 5C and Figure S9E). We noticed that the addition of circRNAs co-translationally somehow lowered the yield of protein, although more proteins were imported into mitochondria with the presence of mecciRNAs (Figure 5B and 5C). Interestingly, when added after the translation of RPA32 protein (post-translation), mecciND1 still increased the mitochondrial importation of RPA32 protein, although with a much less effect compared to the co-translational setup (Figure 5B and 5D). mecciND5 had no effect on the mitochondrial importation of hnRNPA1 protein, when added post-translationally (Figure 5E). These data suggested that mecciRNAs might promote the formation of protein structures favorite for mitochondrial importation of newly synthesized polypeptides; whereas, once the proteins adopted certain structure already, mecciRNAs added post-translationally would have less effect.

**Figure 5.**
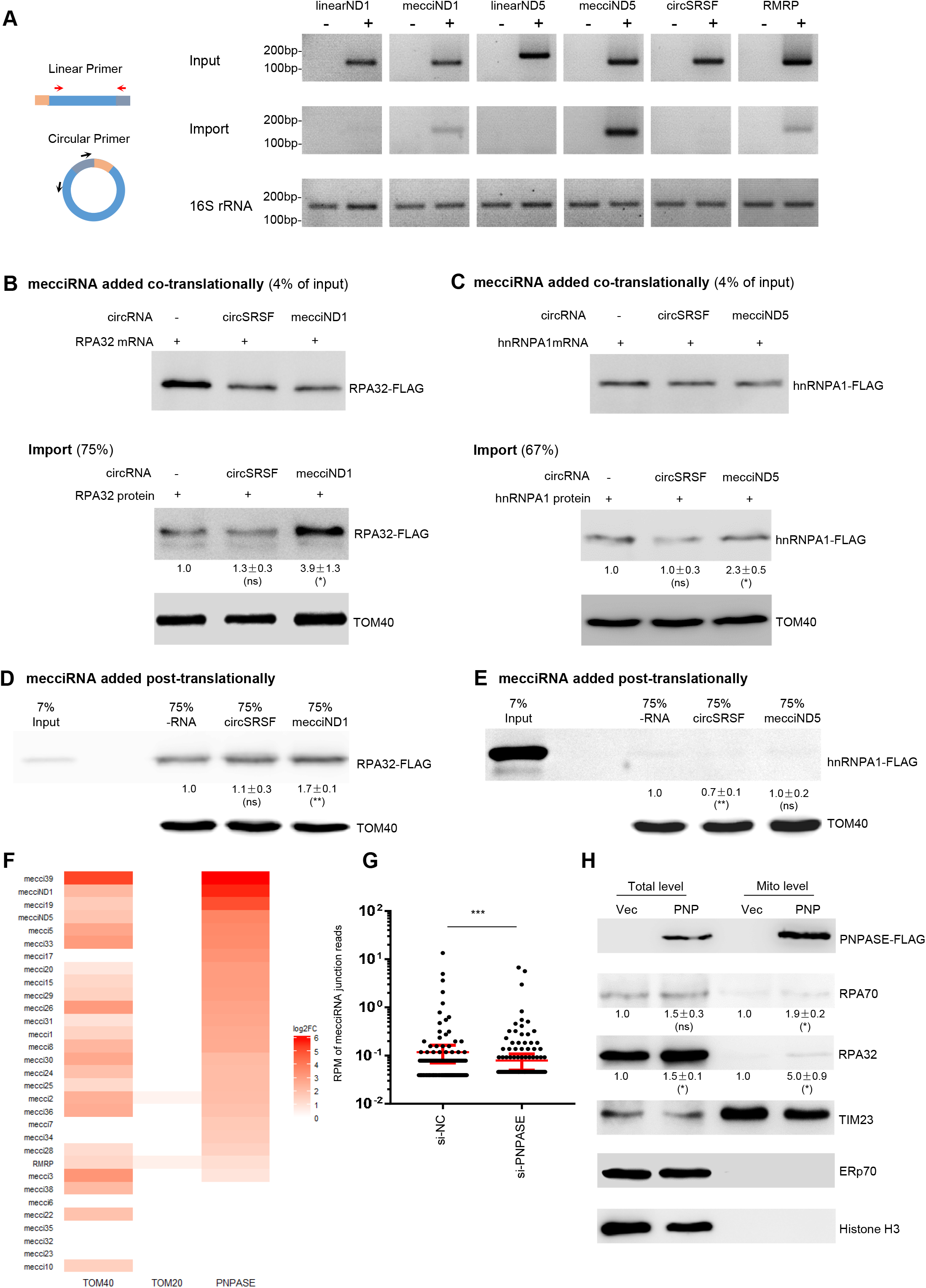
mecciRNAs promote mitochondrial importation in *in vitro* assays and interact with known components of mitochondrial importation. (A) semi-quantitative RT-PCR of *in vitro* RNA import assays. The linear RNA and the counterpart mecciRNA share same sequences. circSRSF, a g-circRNA control; RMRP, a nuclear encoded linear RNA known can be imported into mitochondria, positive control; 16S rRNA, mitochondria encoded rRNA, loading control and a marker for mitochondrial integrity. (B) mecciND1 added together with RPA32 mRNA in the rabbit reticulocyte translational system (co-translation) and the mitochondrial importation of *in vitro* translated RPA32-FLAG protein. Upper, translation products of RPA32-FLAG protein (Input); lower, RPA32-FLAG protein imported into mitochondria. (C) mecciND5 added together with hnRNPA1 mRNA in the rabbit reticulocyte translational system (co-translation) and the mitochondrial importation of *in vitro* translated hnRNPA1-FLAG protein. Upper, translation products of hnRNPA1-FLAG protein (Input); lower, hnRNPA1-FLAG protein imported into mitochondria. (D) mitochondrial importation of *in vitro* translated RPA32-FLAG protein, mecciND1 added post-translationally. (E) mitochondrial importation of *in vitro* translated hnRNPA1-FLAG protein, mecciND5 added post-translationally. For (B)-(E), circSRSF, a g-circRNA control; TOM40, mitochondria outer membrane protein, a loading control; relative importation efficacy of protein was calculated. (F) RNA immunoprecipitation (RIP) against FLAG-tagged TOM40, TOM20 and PNPASE with lysate of transfected 293T cells. Levels of mecciRNA enrichment were examined by Real-time qPCR, and then the fold changes (log2FC) compared to the IgG control were converted into the Heatmap. (G) Mitochondrial mecciRNA levels from mitochondrial RNA-seq data after knockdown of PNPASE. Error bars, s.e.m.; ***P < 0.001 by two-tailed Mann–Whitney U test. (H) Mitochondrial RPA protein levels under the overexpression of PNPASE protein in 293T cells; Vec, vector control; PNP, PNPASE overexpression. TIM23, a mitochondria inner membrane protein, loading control; ERp70, ER marker; Histone H3, nuclear marker. In (B-E) and (H), quantification of proteins levels was normalized to mitochondrial loading controls (TOM40 or TIM23), and the relative levels were the ratios to the corresponding control. n=3 independent experiments; data were mean±s.e.m; ns, not significant; *P < 0.05; **P < 0.01; Student’s *t*-test.

We also found that mecciND1 and mecciND5 imported into mitochondria could be exported out in the *in vitro* assays (Figure S9F). This result was in consistent with the cellular localization of mecciRNAs in both mitochondria and cytosol; mecciRNAs might shuttle between mitochondria and cytosol.

### mecciRNAs interacted with known components of mitochondrial importation

It would be reasonable to propose that mecciRNAs such as mecciND1 and mecciND5 interact with complexes such as TOM40 to facilitate protein entry into mitochondria. RNA-IP against TOM40 complex proteins TOM40 and TOM20 was performed, and indeed mecciRNAs including mecciND1 and mecciND5 were found to interact with TOM40 (the channel-forming protein of TOM40 complex), but not TOM20 (the major import receptor for mitochondrial signal peptide sequences) (Harbauer et al., 2014; Wiedemann and Pfanner, 2017) (Figure 5F and Figure S10A and S10B). PNPASE as a major mitochondrial RNA importation factor was also examined, and most mecciRNAs examined interacted with PNPASE (Figure 5F and Figure S10A, S10B). Knockdown of PNPASE led to decreased mitochondrial mecciRNA levels (Figure 5G and Figure S10C). Overexpression of PNPASE increased mitochondrial RPA protein levels and mecciND1 levels (Figure 5H and Figure S10D). It was known that a small stem-loop structure in RNAs could facilitate mitochondrial RNA importation through PNPASE (Wang et al., 2010). There was a predicted small stem-loop in both mecciND1 and mecciND5, and data from published icSHAPE assay for detecting *in vivo* RNA secondary structure demonstrated that the stem-loop structures in mecciND1 and mecciND5 might be actual (Sun et al., 2019) (Figure S11).

### Dynamic expression of mecciND1 and its association with cellular physiology

Under both UV and hydrogen peroxide treatments, increased mecciND1 levels were observed with a positive correlation to increased mitochondrial RPA70 and RPA32 protein levels (Figure 6A and 6B). Total protein levels of RPA32 were not changed under both conditions, and those of RPA70 were not changed under UV treatment but increased under hydrogen peroxide treatment (Figure 6A and 6B). Levels of mecciND1 also increased upon cellular stresses of hypoxia and tunicamycin treatments (Herst et al., 2017) (Figure 6C and 6D). UV, hydrogen peroxide, hypoxia, and tunicamycin are known to induce DNA damage and thus trigger DNA repair in mitochondria (Herst et al., 2017; Van Houten et al., 2006). Actually, it is believed that mtDNA is more susceptible to oxidative damage than nuclear DNA under regular or stressed physiological conditions, because mtDNA is located in proximity to the ROS-generating respiratory chain (Herst et al., 2017; Van Houten et al., 2006). RPA70 and RPA32 are well known to form complex and to be essential in the repair and duplication of genomic DNA by binding to single-stranded DNA (Zou et al., 2006). Knockdown of RPA70 and RPA32 respectively decreased copy numbers of mitochondrial DNA (mtDNA) (Figure 6E), indicating that RPA proteins were also involved in the duplication of mtDNA. In consistence with changes in mitochondrial levels of RPA proteins, mtDNA copy numbers were decreased upon mecciND1 knockdown and were increased under mecciND1 overexpression (Figure 6F).

**Figure 6.**
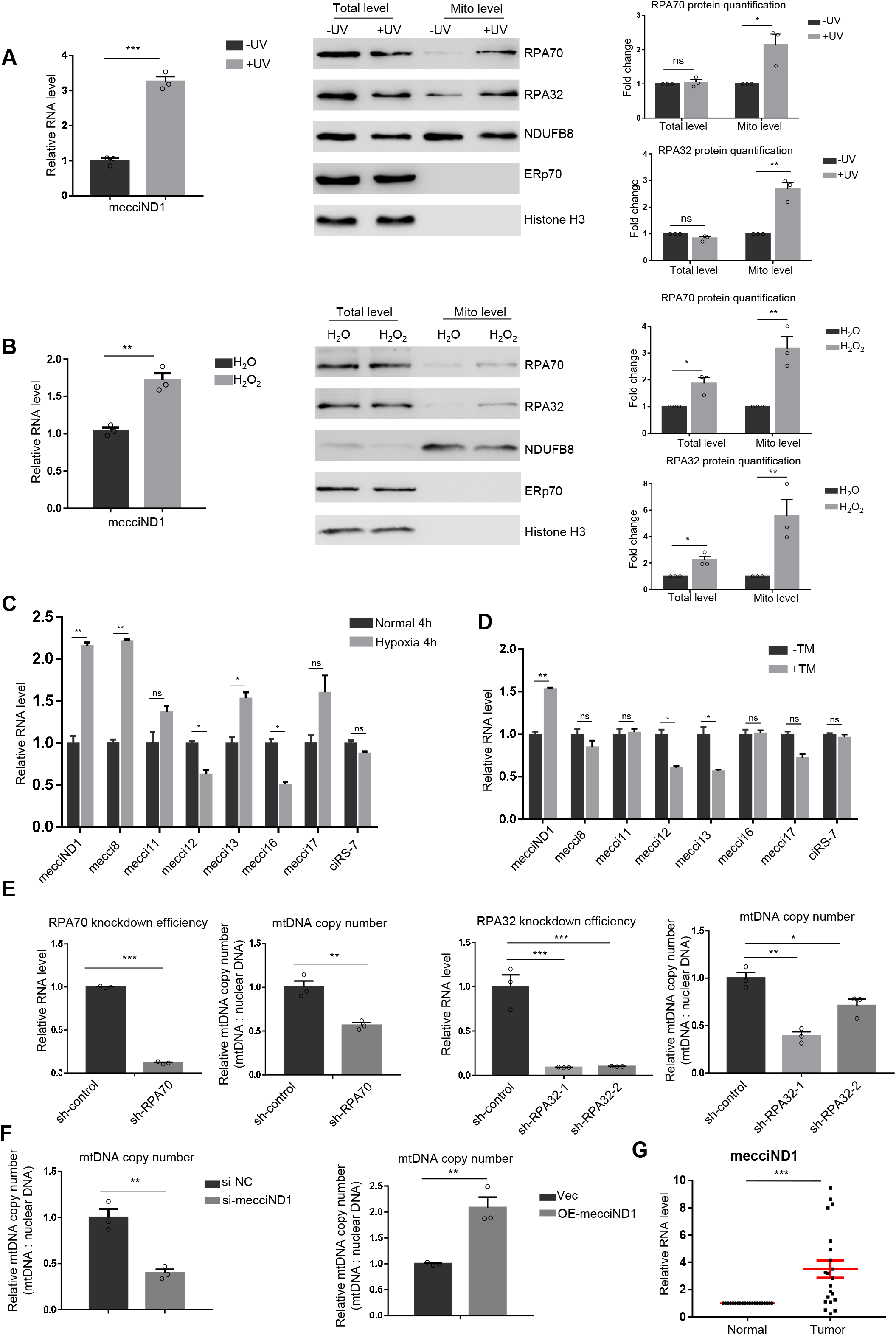
Dynamic expression of mecciND1 and its association with cellular physiology. (A) MecciND1 levels and mitochondrial RPA70 and RPA32 protein levels were increased upon UV irradiation in RPE-1 cells. Quantification of RPA proteins is shown (normalized to NDUFB8). (B) mecciND1 levels and mitochondrial RPA70 and RPA32 protein levels were increased after H_2_O_2_ treatment in RPE-1 cells. Quantification of RPA proteins is shown (normalized to NDUFB8). (C), (D) Expression of mecciND1 increased under 4 h hypoxia culture (C), and 2 h tunicamycin (TM) treatment (D). (E) Knockdown of RPA70 or RPA32 decreased mitochondria DNA (mtDNA) copy number. The knockdown efficiency of RPA70 and RPA32 at mRNA level is also shown; sh-control, shRNA with scrambled sequence. (F) Copy number of mtDNA under mecciND1 knockdown or overexpression. (G) mecciND1 levels in pairs of tumor samples and adjacent tissues from 21 Hepatocellular carcinoma (HCC) patients. In (A)-(E) and (G), qRT-PCR relative RNA levels were normalized to 18S rRNA; in (A)-(F), n=3 independent experiments; in (A)-(G), error bars, s.e.m.; ns, not significant; *P < 0.05; **P < 0.01; ***P < 0.001 by Student’s *t*-test.

Mitochondria are closely related to cancers (Vyas et al., 2016; Zong et al., 2016). We examined mecciND1 levels in pairs of tumor samples and adjacent tissues from 21 Hepatocellular carcinoma (HCC) patients, and mecciND1 was significantly upregulated in HCC (Figure 6G). mecciND5 was also upregulated in HCC, although not as strikingly as mecciND1 (Figure S12A). mecciRNAs corresponding to *S. pombe* mitochondrial genes (*cox1*, *cob1*, *21S rRNA*) and *C. elegans* mitochondrial genes (*nduo-1*, *nduo-5*, and *ctc-1*) were also identified, suggesting that mecciRNAs and their physiological functions might be ubiquitous in eukaryotes (Figure S12B and S12C).

## DISCUSSION

Mutual relations between mitochondria and the “host” cell are fundamental inquiries of biology. Based on results from this study and previous findings, a proposed model in which mecciRNAs facilitate the mitochondrial entry of nuclear encoded proteins through the TOM40 complex is suggested (Figure S12D). mecciRNAs may promote the formation of structures favorite for mitochondrial importation by interacting with nascent polypeptides in the cytosol (Figure 5B and 5C), which is a mechanism of molecular chaperone like previously known for heat shock proteins (Schmidt et al., 2010). Actually, it has been shown that 5S rRNA acts as a molecular chaperone in the co-importation of 5S rRNA and rhodanese, in which 5S rRNA interacts co-translationally with 5S rRNA and leads to an enzymatically inactive conformation for rhodanese in human cells, and then both 5S rRNA and rhodanese get imported into mitochondria (Smirnov et al., 2010).

mecciND1 and mecciND5 interact with TOM40 to facilitate mitochondrial protein entry (Figure 5F), but whether they go through the TOM40 complex together with their protein patterns as a ribonucleoprotein complex remains unknown. It is highly possible that mecciRNAs facilitate mitochondrial entry of proteins with RNA binding ability, since the process involves interaction between mecciRNAs and the targeted proteins. hnRNPA proteins as mecciND5 partners are well-known RNA binding proteins (Geuens et al., 2016). RPA proteins as mecciND1 partners are ssDNA binding proteins with very high affinity (Kd ~10^−10^ M), although they can also bind to RNA with a much lower affinity of ~10^−6^ M (Kd), which is still a reasonable affinity for RNA binding proteins (Brill and Stillman, 1989; Kim et al., 1992). Many so called noncanonical RNA binding proteins bind RNAs transiently or with low affinity may also be protein partners of specific mecciRNAs (Moore et al., 2018). mtSSB is generally regarded as the mitochondrial ssDNA binding protein, although almost no study has been performed to examine whether RPA proteins also play roles in mitochondria. Previous proteomics investigations have demonstrated presence of RPA proteins in mitochondria (Smith and Robinson, 2015), and our data directly show the existence and function of RPA proteins in mammalian mitochondria (Figure 2, 3 and 6). Mitochondria require a substantial number of nucleic acid binding proteins encoded by nuclear genome for their RNA metabolism, transcription, translation, and DNA duplication (Antonicka and Shoubridge, 2015; Calvo et al., 2015; Lopez Sanchez et al., 2011; Zhang et al., 2014). Presumably, most of these proteins would bind to multiple kinds of RNAs in the cytosol, since RNA binding proteins are generally with a broader spectrum of RNA targets or with less specificity. Allowing mecciRNA-bounded proteins to enter the mitochondria may even serve as a sorting mechanism to make sure the entry of right types and amounts of proteins and to keep unwanted cytosolic proteins and RNAs from entering the mitochondria. Interestingly, in a recent study, many proteins are found to demonstrate aberrant mitochondrial targeting in the absence of SRP that facilitates protein targeting to the endoplasmic reticulum (Aviram and Schuldiner, 2017; Costa et al., 2018; Walter and Blobel, 1982). There are also g-circRNAs encoded by the nuclear genome in the mitochondria (Figure 1B), and it is also possible for them to play similar functions like mecciRNAs in mitochondrial protein importation. The fact that both mecciND1 and mecciND5 overexpressed with plasmids can facilitate the mitochondrial importation of their corresponding target proteins somewhat supports this speculation (Figure 3B, 4B and Figure S7C, S8C). It is also possible for mecciRNAs to play roles other than facilitating mitochondrial protein importation. Future researches are in demand to further examine mecciRNAs in mitochondrial protein importation and in other potential roles.

MecciRNAs are distributed both in and outside of the mitochondria. We provide indications that mecciRNAs may shuttle in and out of the mitochondria, but how they are exported and imported remains elusive. mecciRNAs interact with PNPASE, an enzyme has been demonstrated to be essential for mitochondrial importation of RMRP and several other noncoding RNAs (Cheng et al., 2018; Wang et al., 2010). Recently, it is shown that dsRNAs derived from mitochondria play PNPASE related roles (Dhir et al., 2018). The interaction between PNPASE and mecciRNAs is involved in the protein importing role of mecciRNAs (Figure 5F-H). PNPASE may be simply required for the mitochondrial importation and exportation of mecciRNAs, and thus to affect protein importation indirectly. PNPASE plays essential roles in mitochondria, and a recent study shows that knockout of PNPASE gene in cultured cells leads to depletion of mtDNA, and the cells eventually are absent of mitochondria (rho0 cells) (Shimada et al., 2018). The relevance of PNPASE to mecciRNAs and the mitochondrial protein importation extends the understandings about critical roles of PNPASE.

Junction sites as well as the 5’ & 3’ flanking sites of mecciRNAs resemble those of g-circRNAs generated from backsplicing, arguing strongly for a mechanism of backsplicing in the biogenesis of mecciRNAs (Figure 1C and Figure S1B, S1C). There are also recent indications about possible existence of some splicing factors in mammalian mitochondria (Herai et al., 2017). Introns and linear splicing events are present in yeast mitochondria but are generally absent in mitochondria of multicellular animals (Asin-Cayuela and Gustafsson, 2007; Gaspari et al., 2004; Gray et al., 1999, 2001), and linear splicing is absent for mitochondrial transcripts in our RNA-seq data. Nuclear transfected plasmids harboring the corresponding mitochondrial DNA fragment (without any extra sequences such as those complementary flanking sequences known to promote backsplicing) can successfully overexpress mecciRNAs (Figure 3B, 4B and Figure S7C, S8C). It seems that multicellular animals still hold the mitochondrial splicing in the form of backsplicing to generate mecciRNAs, although mitochondria may lack essential components of linear splicing; the missing can be due to lacking of real intron on mitochondrial pre-RNA transcripts, or simply due to absence of protein factors required. Biogenesis and metabolism of mecciRNAs require further elucidation. There are also examples that circRNAs generated from backsplicing are identified for single exon genes of the nuclear genome, and no linear splicing is involved in the biogenesis of mRNA from single exon genes (Glažar et al., 2014); for example, the well-known circSry in mice is encoded by a single exon gene Sry (Capel et al., 1993; Hansen et al., 2013; Memczak et al., 2013).

Mitochondria need to maintain their homeostasis and react to different physiological conditions. Previous researches have focused on how cells “manage” mitochondrial protein importation in response to stresses (Harbauer et al., 2014; Quirós et al., 2015; Sokol et al., 2014; Weidberg and Amon, 2018). It has just recently been shown that a mitochondria initiated surveillance pathway mitoCPR (mitochondrial compromised protein import response) can mediate the degradation of unimported proteins from the mitochondrial surface in the budding yeast (Weidberg and Amon, 2018). The identification of mitochondrial encoded mecciRNAs and their potential physiological roles associated with stress responses and cancer expand the complexity of mitochondrial RNAomics as well as categories and functions of circRNAs (Chen, 2016; Chen et al., 2015; Hansen et al., 2013; Li et al., 2015; Memczak et al., 2013; Mercer et al., 2011; Salzman et al., 2012; Szabo and Salzman, 2016; Zhang et al., 2014), and mecciRNAs are another layer of regulators of mitochondrial physiology and mitochondria-to-nucleus relations.

## Supporting information

All the supplemental information

## SUPPLEMENTAL INFORMATION

includes full detailed methods, 12 Figures, and 1 Table.

## AUTHOR CONTRIBUTIONS

G. S. conceived the study, supervised the project, and provided funding. X. L. performed the majority of the experiments, analyzed data and together with G. S. wrote the manuscript. X. W. and J. L. contributed to the bioinformatics analysis. S. H., Y. D., and H. Y. performed some of the experiments. X. B., Q. Z., and G. W. provided consulting. B. W. provided experimental materials. Q. S. provided SIM platform and technical support.

## ACKNOWLEDGEMENTS

Thanks for Lei Liu in technical support of *C. elegans* experiments. This work was supported by grants to G.S.: the National Basic Research Program of China (2015CB943000), the National Key R&D Program of China (2018YFC1004500), the National Natural Science Foundation of China (31725016 and 31600657), and the Strategic Priority Research Program (Pilot study) “Biological basis of aging and therapeutic strategies” of the Chinese Academy of Sciences (XDPB10).

