## Supplementary material for "The identification of mecciRNAs and their roles in mitochondrial entry of proteins": All the supplemental information

**Supplemental Information of full Methods and figure legends of Supplemental Figures**

#### MATERIALS AND METHODS

##### Cell cultures and model organisms

HeLa, HEK293T, RPE-1, HepG2, N2a and NIH3T3 cells were all originated from the ATCC and cultured with DMEM. All cells were cultured under standard conditions including 10% FBS and 1% penicillin/streptomycin at 37 °C under 5% $CO_2$. Cells were tested for mycoplasma by DAPI staining, to ensure the absence of contamination. *S. pombe* (PR109), N2 wild-type *C. elegans* strain, *D. rerio* (AB, adult male) and 6-week-old male C57BL/6 wild-type mice were cultured according to standard methods. Protocols involving mice and *D. rerio* were approved by the Institutional Animal Care and Use Committee at the University of Science and Technology of China.

##### Clinical Samples

All fresh HCC patient tumor samples and adjacent tissues were collected from The First Affiliated Hospital of University of Science and Technology of China, which was approved by the Human Research Ethics Committee of University of Science and Technology of China (USTCEC201700007). Written informed consent was obtained from each patient for this study. All samples were rinsed with DEPC water and then kept in RNAhold (Transgene) within 30 minutes after removing from the operation. HCC patient tumor sample and adjacent tissue pairs were collected from 21 patients (12 males and 9 females with advanced stage HCC, all of them were HBsAg positive, and did not have anti-tumor therapy before surgery) (Sheng et al., 2019).

##### DNA & RNA isolation, RNase R treatment, and PCR reactions

Genomic and mitochondrial DNA were extracted by RNase A and proteinase K digestion, phenol and chloroform extraction with isopropanol precipitation. Total RNA was extracted with TRIzol Reagent (Invitrogen) according to the manufacturer's procedures. For RNase R treatment, 5 μg total RNA was treated with 8 U RNase R (Epicentre) in 50 μl total volume for 40 min. Complementary DNA (cDNA) was synthesized from RNA with the GoScript Reverse Transcription System (Promega) according to the supplied protocol, with random hexamer primers. Quantitative PCR (qPCR) was performed with GoTaq SYBR Green qPCR Master Mix (Promega) on a PikoReal 96 real-time PCR system (Thermo Scientific) according to standard procedures. For semi-quantitative PCR, 18-20 amplification cycles were performed.

##### RNA sequencing

For high-throughput sequencing, RNAs isolated from mitochondria were iron-fragmented at 95℃ and then subjected to end repair and 5'-adaptor ligations. Then, reverse transcription was performed with random primers containing 3'-adaptor sequences and randomized hexamers. After cDNA purification and amplification, the PCR products of 200–500 bp were purified and quantified. Libraries were prepared according to the manufacturer's instructions and subjected to 151-nt paired-end sequencing with an Illumina Nextseq 500 system. We sequenced each library to a depth of 10–50 million read pairs and then removed adapters with cutadapt to obtain clean reads.

**DNA re-sequencing**

The extracted genomic or mitochondrial DNA was sequenced using the HiSeq 2500 sequencing systems (Illumina Inc., https://www.illumina.com). In short, the obtained reads were mapped to hg19 reference genome with bowtie (-v 1) and PCR amplifications were performed with PICARD.

**MecciRNA identification and g-circRNA identification**

For mecciRNA annotation, the pipeline of find_circ (Memczak et al., 2013) was applied with reference genome (human: hg19; mouse: mm9; zebra: danRer11) downloaded from the UCSC genome browser (http://genome.ucsc.edu/). In brief, we aligned reads to the reference genome and filtered reads that aligned contiguously; for the remaining reads, we extracted 20-mers from both ends and aligned them independently to find unique anchor positions in chrM (mitochondrial genome), and then we extended the anchor alignments of two read segments mapping to chrM in the reversed order. Only circRNAs with ≥2 junction reads across four human samples or four mouse samples were analyzed. For zebrafish with only one sample sequenced, circRNAs with ≥1 junction reads were analysed. g-circRNAs (circRNAs encoded by the nuclear genome) were identified with find_circ.

**Motif identification**

The enriched motifs of junctions (-len 10) and flanking site sequences (-len 20) for mecciRNAs were generated by HOMER.

**Linear splicing identification in mitochondrial transcripts**

For linear splicing, we first aligned the RNA-seq data to the reference genome with bowtie (-v 1) to filter the contiguously mapped reads. Then, we mapped the remaining reads to chrM with blat to find the *de novo* split transcripts in mitochondrial.

**Northern blotting**

Sense and antisense digoxigenin-labeled RNA probes were prepared with a DIG Northern Starter Kit (Roche) according to the manufacturer’s protocol. 20 µg of total RNA with or without RNase R digestion and RiboRuler Low Range RNA Ladder (Thermo Scientific) were loaded in 8% Urea PAGE gel and run for 1 h in 1X TBE buffer. RNA was transferred onto Hybond-N+ membranes (GE Healthcare) by electronic transfer. After transfer, the membranes were UV-crosslinked and hybridized with specific RNA probes according to the manufacturer’s protocol (Roche, DIG Northern Starter Kit). Images were taken with an ImageQuant LAS4000 Biomolecular Imager (GE Healthcare) (Wang and Shan, 2018).

**Plasmid construction**

All plasmids were constructed with restriction-enzyme digestion (Thermo Scientific FD) and ligation (Promega A3600) or with recombinant methods (Vazyme c112-02). Oligonucleotide sequences for primers used in plasmid construction, probe preparation, siRNAs, and biotin-labeled nucleic acids are listed in Table S1. The shRNA plasmid for knockdown of hRPA70 mRNA (shRPA70, TRCN0000010985), hRPA32 mRNA (shRPA32-1, TRCN0000005986; shRPA32-2, TRCN0000005987) with negative-control shRNA (shC002) was obtained from the MISSION shRNA Library (Sigma-Aldrich). For the overexpression of C-terminal FLAG-tagged TOM20, TOM40 and PNPASE the plasmid backbone was pmR-mCherry. The structures of overexpression plasmids of human mecciND1 and mecciND5 are shown in Figure S7C and Figure S8C. The major constructions of plasmids used for *in vitro* assays were shown in Figure S9B (Wesselhoeft et al., 2018). Purified mecciND1, mecciND5, circSRSF, and RMRP fragments from PCR reactions and chemically synthesize group I self-splicing intron were cloned into pUC57 vector. Coding sequence of RPA32 or hnRNPA1 fused with sequence corresponding to FLAG tag at 3' terminal was inserted downstream of the SP6 promoter in pcDNA3 plasmid. All plasmids were sequenced for confirmation.

**RNA pull-down with biotin-labeled antisense oligonucleotides**

RNA pull-down with 5'-biotinylated antisense (AS) oligos was modified with a previously described method (Hu et al., 2016). Briefly, Cells were cross-linked for 2 min in a UV cross-linker (UVP) at 120 mJ/cm$^2$ strength. The cross-linked cells or purified mitochondria were then lysed in RIPA buffer (50 mM Tris-HCl, pH 8.0, 150 mM NaCl, 5 mM EDTA, 1% NP-40, 0.1% SDS), 2 mM DTT, 1X protease-inhibitor cocktail (Roche) and 200 units/ml RNasin® Ribonuclease Inhibitors (Promega) for 10 min on ice, then sonicated for 10 min with a Sonics Vibra-Cell. Lysates were cleared of cell debris by centrifugation at 13,000 g for 15 min. 100 pmol biotinylated AS oligos were added to the supernatant and mixed by end-to-end rotation at room temperature for 2 h. M-280 Streptavidin Dynabeads (Life Technologies) were blocked with 500 ng/μl yeast tRNA and 500 ng/μl BSA for 1 h at room temperature, then washed with RIPA buffer before being resuspended. 50 μl blocked Dynabeads was added per 100 pmol of biotin-DNA oligonucleotides, and the mixture was then rotated for 2 h at room temperature. Beads were captured with magnets (Life Technologies) and washed two times with RIPA buffer supplemented with 500 mM NaCl. RNAs and proteins were eluted from beads for further analysis. Proteins pulled down by AS oligos were separated on SDS–PAGE gels, and silver-stained.

**Mass spectrometry**

Specific silver-stained bands were cut, digested and extracted. The masses of the peptides in the extract were then measured by MS to obtain the peptide mass fingerprints. Next, peptides were selected to undergo fragmentation via tandem MS. Both the MS and tandem MS data were searched against protein sequence databases to determine the proteins present in the gel.

**RNA Immunoprecipitation (RIP)**

RIP was carried out as previously described with some modifications (Li et al., 2015). Briefly, Cells were cross-linked in a UV cross-linker (UVP) at 120 mJ/cm$^2$ strength. and then harvested in ice-cold RIPA buffer (50 mM Tris-HCl, pH 8.0, 150 mM NaCl, 5 mM EDTA, 1% NP-40, 0.1% SDS), 200 units/ml RNasin® Ribonuclease Inhibitors (Promega), 2 mM DTT, and 1X protease-inhibitor cocktail (Roche). Cells were sonicated for 10 min with a Sonics Vibra-Cell, the cell suspension was centrifuged at 13,000 g for 15 min at 4 °C, and the supernatant was collected. Antibody or IgG (as control) was added and incubated 4 h at 4 °C for antigen coupling, and then Protein G Dynabeads (Life Technology) suspension was then added and allowed to bind for at least 2 h at 4 °C. The Protein-antibody–beads complexes were washed two times with RIPA buffer and two times with RIPA buffer supplemented with 500 mM NaCl. One-fifth of the beads after the last wash was heated 100 °C for 10 min in SDS-loading buffer and then saved for Western blotting. The remaining Protein-antibody–beads complexes were digested with proteinase K at 55 °C for 30 min, followed by extraction with TRIzol Reagent to obtain RNA. The following antibodies were used: anti-RPA70 (Abcam, ab79398); anti-RPA32 (Abcam, ab2175); anti-hnRNPA1 (Sigma-Aldrich, R4528); anti-FLAG (Sigma-Aldrich, F1804). Antibody validation is provided on the manufacturers' website.

**Western blotting**

For Western blotting, whole cell lysates, mitochondria lysates, and IP mixtures were separated on SDS–PAGE gels and then transferred to BioTrace NT Nitrocellulose Transfer Membrane (PALL Co.). Membranes were processed according to the ECL Western Blotting protocol (GE Healthcare). The following antibodies were used in Western blotting: anti-RPA70 (Abcam, ab79398); anti-RPA32 (Abcam, ab2175); anti-hnRNPA1 (Sigma-Aldrich, R4528); anti-hnRNPA2B1 (Abcam, ab6102), anti-hnRNPA3 (Proteintech Group, 25142-1-AP); anti-GAPDH (Proteintech Group, 60004-1-IG); anti-ERp70 (Proteintech Group, 14712-1-AP); anti-NDUFB8 (Proteintech Group, 14794-1-AP); anti-TOM20 (Proteintech Group, 11802-1-AP), anti-TOM40 (Proteintech Group, 18409-1-AP); anti-TIM23 (Proteintech Group, 11123-1-AP); anti-FLAG (Sigma-Aldrich, F1804); anti-β-actin (Transgene, HC201); Anti-PNPT1 (Abacm, ab96176). Antibody validation is provided on the manufacturer' s website. Quantification of Western blot bands was performed with Image J.

**Immunofluorescence (IF) combined with fluorescence in situ hybridization (FISH)**

FISH probes were generated with Transcript Aid T7 High Yield Transcription Kit (Thermo Scientific), and then labeled with Alexa Fluor546, 488 or 647, by using the ULYSIS Nucleic Acid Labeling Kit (Invitrogen), which added a fluor on every G in the probe to amplify the fluorescence intensity. HeLa cells or RPE-1 cells were grown on coverslips, fixed in 3% PFA for 10 min at RT, and then permeabilized with freshly made PBS, 1% v/v Triton X-100 (plus 200 units/μl RNase inhibitor) on ice for 10 min. Blocked in 1% w/v BSA for 30 min at RT. Incubated with primary antibody 1:100 diluted in 1% BSA (containing 200 units/ml RNase inhibitor) for 3 h at RT, washed with PBS, 0.1% Triton X-100 for three times. Incubated with secondary antibody (1:200 diluted in 1% BSA, containing 200 units/ml RNase inhibitor) for 1 h at RT in a dark and humid chamber (made with paper tissues soaked with PBS), washed with PBS, 0.1% Triton X-100 for three times (dark), and washed with 2X SSC (Sigma-Aldrich) once (dark). RNA probes were denatured at 80 °C for 10 min with 30 ng/μl Salmon DNA (Invitrogen) and 500 ng/μl yeast tRNA (Invitrogen) in 2X hybridization buffer (4X SSC, 40% w/v dextran sulfate). After denature, add 200 units/ml RNase inhibitor in the mix and put on the slides. Hybridize overnight at 37 °C in a dark and humid chamber (made using paper tissues soaked in 50% v/v formamide in 2X SSC). Slides were washed with 2X SSC 0.1% Triton X-100 at 45 °C; counterstain DNA with DAPI. Cover-slide was then mounted. The following antibodies were used in IF: anti-RPA70 (Abcam, ab79398); anti-RPA32 (Abcam, ab2175); anti-hnRNPA1 (Sigma-Aldrich, R4528); anti-hnRNPA2B1 (Abcam, ab6102), anti-hnRNPA3 (Proteintech Group, 25142-1-AP); anti-TOM20 (Proteintech Group, 11802-1-AP; Abcam, ab56783), anti-TOM40 (Proteintech Group, 18409-1-AP). Donkey anti-Mouse Secondary Antibody, Alexa Fluor 488 (Invitrogen, A21202), Donkey anti-Rabbit IgG Secondary Antibody, Alexa Fluor 546 (Invitrogen, A10040), Goat Anti-Rabbit Secondary Antibody Alexa Fluor® 488 (Abcam, ab181448), Goat Anti-Mouse Secondary Antibody Chromeo™ 546 (Abcam, ab60316), Antibody validation is provided on the manufacturers' websites.

**Transfection of plasmids, siRNAs and AMOs**

Plasmid and siRNA transfection were conducted with Lipofectamine 2000 (Invitrogen) according to the supplier's protocols. All siRNAs were subjected to BLAST search to ensure the absence of hits with more than 17-nt matches in the corresponding genomes. Antisense Morpholino oligonucleotides (AMOs), including mecciND1 AMO, mecciND5 AMO, and scrambled AMO, were synthesized at Gene Tools. AMOs were transfected through electroporation with the Nucleofector$^{TM}$ system (Lonza) according to the manufacturer's instructions. Cells were harvested for analysis or downstream experiments 8-12 h after AMO transfection. The final concentration of AMOs was 10 μM.

**Quantification of mecciRNA copy number per cell**

DNA fragments corresponding to human mecciND1 and mecciND5 were amplified through RT-PCR reactions with divergent primers (to amplify mecciRNA only). The purified DNA fragments were diluted by grads multiple to plot standard curves through real-time PCR. Total RNAs from 1.0 X $10^6$ HeLa, 293T, RPE-1 and HepG2 cells were extracted, and cDNAs were then synthesized. The mecciND1 and mecciND5 copy numbers per cell in each cell line were calculated on the basis of cell numbers and the Ct values from the qRT-PCR using the standard curves.

**UV, H2O2, TM, and hypoxia treatment of cells**

For UV irradiation, cells were first grown to reach 80%–90% confluency, then culture medium was discarded, cells were either left untreated (negative control) or exposed to 25 mJ/cm$^2$ UVC irradiation. Cells were then washed with PBS and placed in fresh media, and then incubated for 30-60 min before harvest. For $H_2O_2$ treatment, cells were first grown to reach 80%–90% confluency in complete medium and the culture medium was subsequently replaced with DMEM without FBS, together with or without 1 mM $H_2O_2$, cells were then incubated for 1 h before harvest. For TM (tunicamycin) treatment, cells were treated with 100 mg/ml TM for 4 h. Cells exposed to hypoxia were maintained at 1% $O_2$/5% $CO_2$/balance $N_2$ at 37 °C in a modular incubator chamber for 24 h.

**Mitochondria isolation**

Mitochondria isolation of culture cell was carried out as previously described with some modifications (Wieckowski et al., 2009; Williamson et al., 2015). Briefly, cells were harvested and washed by PBS, collected by centrifugation and re-suspended in ice-cold isolation buffer 1 (225 mM mannitol, 75 mM sucrose, 0.1 mM EGTA and 20 mM HEPES-KOH pH 7.4). Cells were homogenized by 20 strokes in a Dounce homogenizer (Kontes). Cell homogenates were centrifuged twice at 1500 g 4 °C for 5 min to discard nuclear and collect clear supernatant. Mitochondria were sedimented at 12,000 g 4 °C for 10 min. The ER contaminant proteins make a large, loose white ring around the more stable mitochondrial pellet. The mitochondrial pellet appeared more yellow in color. This was then gently washed with isolation buffer 2 (225 mM mannitol, 75 mM sucrose, and 20 mM HEPES-KOH pH 7.4) until the white ring was washed out. For mitochondria isolation of animal tissues was carried out as previously described with some modifications (Wieckowski et al., 2009). Briefly, tissues were washed in ice-cold IB-1 (225 mM mannitol, 75 mM sucrose, 0.5% BSA, 0.5 mM EGTA and 20 mM HEPES-KOH pH 7.4); cut into small pieces using scissors in ice-cold IB-3 (225 mM mannitol, 75 mM sucrose and 20 mM HEPES-KOH pH 7.4) and washed once again with fresh 10 ml of ice-cold IB-1. The tissue fragments were resuspended with IB-1 in the ratio 4 ml of buffer per gram of tissue; transferred to the glass tissue homogenizer for Homogenization and then the homogenates were again homogenized in a Dounce homogenizer (Kontes) by 20 strokes. The centrifugation procedures were same as that described in the procedure for mitochondria isolation of culture cell. Zebrafish mitochondria isolation is similar to that for tissue. Briefly, the zebrafish was put on ice for 1 min, washed in ice-cold IB-1, cut into small pieces using scissors in ice-cold IB-3. The follow procedures were same as described in the procedure for mitochondria isolation of culture cell.

**Isolation of mitochondrial RNAs & proteins and cytosolic RNAs**

The purified mitochondria for RNA-seq were then treated with 100 mg/ml RNase A for 10 min on ice following 1 mg/ml Digitonin (Sigma) for another 10 min to remove contaminating cytoplasmic RNA. The purified mitochondria for Western blotting were treated with 1 mg/ml Digitonin (Sigma) for 10 min to remove contaminating protein. Both Western blotting and RT-qPCR were carried out to confirm quality of purified mitochondria. For cytosolic RNA isolation, the supernatants after precipitation of mitochondria was centrifuged again at 15000 g 4°C for 10min followed by extraction with TRIzol LS Reagent to obtain RNA.

***In vitro* transcription and circularization**

Templets of *in vitro* transcription were amplified by specific primers containing T7 RNA polymerase promoter sequences from the vectors described above. Linear RNA or circRNA precursor were synthesized by *in vitro* transcription using a TranscriptAid T7 High Yield Transcription Kit (Thermo Scientific). After *in vitro* transcription, reactions were treated with DNase I for 20min. After DNase treatment, linear RNA was purified using Phenol-chloroform PH ~4.5. For circRNA, the procedures of *in vitro* circularization were shown in Figure S9B and circularization was carried out as previously described with some modifications (Wesselhoeft et al., 2018). Briefly, after DNase treatment, additional GTP was added to a final concentration of 2 mM along with a circularization buffer including magnesium (15 mM $MgCl_2$, 1 mM DTT 50 mM Tris-HCl, PH 7.5) and then the reaction was heated at 55 °C for 20 min. RNA was then purified using Phenol-chloroform PH ~4.5. To enrich for circRNA, 10 μg RNA was digested with RNase R and then Phenol-chloroform purified. RNase R digested RNA was separated on 5% Urea PAGE gel. Bands corresponding to circRNA were excised from the gel and eluted overnight in elution buffer (20 mM Tris-HCl, PH 7.5, 250 mM NaOAc, 1 mM EDTA, 0.25% SDS). After elution, the gel fragments were discarded by centrifugation at 15000g 4 °C for 2 min. RNA was then purified using Phenol-chloroform PH ~4.5. As shown in in Figure S9B, the circRNAs (including the circRNA control) *in vitro* synthesized could be examined from the endogenous circRNAs due to a small stretch of added sequences (29 nt).

**Importation of RNAs into isolated mitochondria**

Mitochondria were isolated from 293T cells. The isolated mitochondria were resuspended in import buffer (250 mM Sucrose 20 mM HEPES-KOH, PH 7.4, 5 mM $MgCl_2$, 60 mM KCl, 2 mM ATP, 10 mM Succinate, 1 mM DTT). After incubation for 15 min at 30°C, the mitochondria (~25-50 μg of total mitochondrial proteins) were divided equally into import system. 200 ng Linear RNA or circular RNA with import buffer were added into the system to the final volume of 100 μl. The import reaction was performed at 30°C for 30 min with shaking gently several times. After the reaction, 3500 gel units Micrococcal Nuclease (NEB) together with its reaction buffer were added to eliminate RNAs that not imported into mitochondria, and then the mixture was incubated at 30°C for 20 min. Mitochondria were then spun down at 13000 g 4°C for 5 min. After 2 times wash with isolation buffer 2, the mitochondrial pellets were dissolved in TRIzol Reagent for RNA.

***In vitro* mitochondrial importation of proteins**

Capped RPA32-FLAG and hnRNPA1-FLAG mRNAs were *in vitro* transcribed from linearized SP6 plasmids using the mMESSAGE mMACHINE™ SP6 Transcription Kit (Invitrogen) according to manufacturer's directions. After transcription, *E. coli* poly(A) polymerase (NEB) was added into reaction for poly(A) tailing of capped-mRNA. For *in vitro* importation of RPA32-FLAG protein together with mecciND1 or hnRNPA1-FLAG protein together with mecciND5 (co-translation), isolated mitochondria were resuspended in import buffer and incubated for 15 min at 30°C. 200 ng circularized mecciND1, mecciND5, circSRSF (g-circRNA control), or no circRNA control along with 2μg RPA32-FLAG or hnRNPA1-FLAG mRNA were added into Rabbit Reticulocyte Lysate System (Promega) for one hour. The mitochondria (~25-50 μg of total mitochondrial proteins) were divided equally into the translation system to a final volume of 50 μl, at 30°C for 1 h. After the reaction, 5μl of the mixture was taken out as input. 3500 gel units Micrococcal Nuclease (NEB) together with its reaction buffer were added and the mixture, and then incubated at 30 °C for 20 min to digest all RNAs not in the mitochondria. Mitochondria were then spun down at 13000 g 4 °C for 5 min and resuspended with 100 μl import buffer. The samples were treated with 25 μg/ml Trypsin for 5 min at room temperature to digest any protein outside of the mitochondria, and then Trypsin was stopped by 1 mM TLCK (Sigma) and Soybean Trypsin Inhibitor (BI). Mitochondria were spun down again at 13000 g 4 °C for 5 min. After 2 times wash with isolation buffer 2, 1/4 of the mitochondrial pellets was taken out for RNA isolation and the left was subjected to Western blotting. For the "post-translation" effects of mecciRNAs, 2 μg RPA32-FLAG or hnRNPA1-FLAG mRNA were added into Rabbit Reticulocyte Lysate System (Promega) in a total volume of 50 μl for 30 min at 30°C, and then 5 μl of the mixture was taken out as input. The remaining materials were incubated with 200 ng circularized mecciND1, mecciND5, circSRSF (g-circRNA control), or no circRNA control for 15 mins at 30 °C, following the addition of isolated mitochondria (pre-warmed in import buffer for 15 min at 30 °C). The mitochondria (~25-50 μg of total mitochondrial proteins) were divided equally into the three reactions to a final volume of 100 μl, 30 °C, 1 h.

**RNA export assay**

RNA export assay was carried out as previously described with some modifications (Wang et al., 2010). After the assays of mecciND1, mecciND5 or circSRSF RNA importation into isolated mitochondria, the mitochondria were then subjected to Micrococcal Nuclease treatment of 3500 gel units Micrococcal Nuclease (NEB) together with its reaction buffer, 30°C for 20 min. 10 mM EGTA was then added to inactivate the Micrococcal Nuclease, and the mitochondria were spun down at 13000 g 4°C for 10 min. The supernatant was removed, and the mitochondrial pellet was washed with 1ml pre-warmed (30°C) isolation buffer 2. The pellet was then resuspended in 160 μl pre-warmed (30°C) simpler import buffer (250 mM Sucrose 20 mM HEPES-KOH, PH 7.4, 2 mM ATP, 10 mM Succinate). For 0 min sample (before the export assay), 80 μl mixture was taken out and spun at 13000 g 4°C for 10 min, and both the supernatant and pellet were kept for RNA isolation. 20 μl Rabbit Reticulocyte Lysate was added into the 80 μl mitochondria in the simpler import buffer, and the export reaction was performed at 30°C for 20 min with gentle shakes several times. After export, the reaction system was spun at 13000g 4°C for 10 min. Both the supernatant and pellet were then subjected to RNA isolation.

**Mitochondria DNA copy number**

The relative mtDNA copy number was calculated as a ratio of mtDNA/nuclear DNA according to a previous study (Venegas and Halberg, 2012). Briefly, cells were lysed in the RIPA buffer and DNA extracted with phenol/chloroform followed by ethanol precipitation. For quantification of mtDNA, a pair of primers (mtDNA forward and mtDNA reverse) that target the tRNA-Leu (UUR) gene were used for qPCR. To quantify nuclear DNA, we used a pair of primers (nucDNA forward and nucDNA reverse) that target the nuclear β2-microglobulin gene for qPCR.

**Confocal microscopy**

IF-FISH images were taken on a Zeiss LSM 880 confocal microscope with 63X 1.40 NA oil-immersion objective, z-stack images were acquired with a resolution of 1024x1024 using ZEN Black confocal software (Zeiss). The images were saved as .czi files and converted to .tiff files using ZEISS ZEN microscope software. Each .tiff file was processed in the open-source image processing software, (Fiji is Just) Image J. The ROI was drawn as a small rectangle and colocalization percentage of proteins and RNAs in 20 ROIs was measured using image J plugin Coloc2. Z project was processed by Image J.

**Structured illumination microscopy (N-SIM)**

Structured illumination microscopy (SIM) super-resolution images were taken on a Nikon N-SIM system with a 100X oil immersion objective lens, 1.49 NA (Nikon). Images were captured using Nikon NIS-Elements and reconstructed using slice reconstruction in NIS-elements. Images of fixed cells for 3D N-SIM were taken using Z -stacks with step sizes of 0.2 μm. The images were saved as .tiff files. Z project was processed by Image J. Fluorescence quantification of mecciND1, RPA32, mecciND5 and hnRNPA1 in RPE-1 cells was processed by Image J. Briefly, we selected a mitochondrion with a square and measured FISH signal (M, magenta) and protein signal (G, green) separately in this square. 30 (for RPA32 and meeciND1) and 39 (for hnRNPA1 and mecciND5) mitochondrial areas were randomly selected for fluorescence quantification.

**Statistical analysis**

Either Student's *t*-tests or Mann–Whitney U tests were used to calculate P values, as indicated in the figure legends. For Student's *t*-tests, the values reported in the graphs represent averages of three independent experiments, with error bars showing s.e.m. After analysis of variance with F tests, the statistical significance and P values were evaluated with Student's *t*-tests. Statistical methods are also indicated in the figure legends.

**DATA AND SOFTWARE AVAILABILITY**

All RNA-seq and DNA resequencing data will be deposited in the NCBI with accession numbers available.

**Figure Legends of Supplemental Figures**

**Figure S1. Features of mecciRNAs, Related to Figure 1**

(A) Maps of mecciRNA junction positions on mitochondrial genome; backsplicing sites are connected by lines. Mitochondria encoded genes are shown on the circle; the outer ring, heavy strand genes; the inner ring, light strand genes. Maps constructed using Circos (Krzywinski et al., 2009). (B) Junction motif of human, mouse and zebrafish g-circRNAs. (C) Motif for 5' and 3' flanking sites of human, mouse and zebrafish mecciRNAs. (D) Distance between flanking sites of mecciRNAs.

**Figure S2. Experimental verification of mecciRNAs, Related to Figure 1**

(A) Northern blots of mecciND1 and mecciCYB (sense probe, as a negative control). (B) divergent and convergent primers design was shown. (C) PCR with divergent and convergent primers for the verification of mecciRNAs. GAPDH mRNA, negative control; g+mtDNA, nuclear and mitochondrial DNA; RT, reverse transcription; for several gel images, a small open triangle is used to indicate the specific band from mecciRNA. (D) Real-time qPCR and regular PCR showing the resistance of human mecciRNAs to RNase R digestion in HeLa cell. ciRS-7, positive controls; GAPDH mRNA, negative control. (E) Real-time qPCR and regular PCR showing resistance of mouse mecciRNAs to RNase R digestion in HeLa cell and N2a cell. GAPDH mRNA, negative control; R+, R-, with or without RNase R digestion. In (D) and (E), error bars, s.e.m.; n=3 independent experiments.

**Figure S3. Analyses of related NGS data, Related to Figure 1**

**(A)** No DNA read matching junction sequences of human mecciRNAs in re-sequencing reads of genomic DNA (Nuc-DNA) and mitochondrial DNA (Mt-DNA) from HeLa cells. (B) In RNA-seq data of cells without mitochondria (Rho0 MEF cells), no mecciRNA was identified, whereas mecciRNAs were found in wildtype MEF cells (Shimada et al., 2018). (C) Nascent circRNAs in HeLa cells identified from RNA-seq data (Bao et al., 2018).

**Figure S4. Mitochondrial and cytosolic distributions of mecciRNAs, Related to Figure 1**

(A) g-circRNAs and mecciRNAs from sequencing data of cytoplasmic (cyto) and mitochondrial (mito) RNAs (HeLa cells). (B) mecciRNAs have both mitochondrial and cytosolic distributions. ATPase6 is a mitochondrial encoded mRNA. The mitochondrial 16S rRNA is used as endogenous control in qRT-PCR. (C) IF of TOM20 together with FISH signals of ND1 mRNA and mecciND1. Boxed areas are enlarged. Colocalization between ND1 FISH signals (G, green) and TOM20 (R, red) is shown (n=20 randomly selected areas). R/G, The proportion of red signal to green signal colocalization; G/R, The proportion of green signal to red signal colocalization. In (C), scale bars, 5 μm and 500 nm (enlarged areas); in (B), error bars, s.e.m.; *P < 0.05; **P < 0.01; ***P < 0.001 by Student's *t*-test.

**Figure S5. mecciND1 interacts with RPA70 and RPA32, Related to Figure 2**

(A) The position of antisense oligo and mecciND1 pulldown efficiency are shown; Actin mRNA served as negative control. (B) Pulldown of mecciND1 with biotin labeled antisense oligo (AS oligo) in HeLa mitochondria lysate. RPA70 and RPA32 co-pulled down with mecciND1 were verified by Western blots; NDUFB8, negative control. (C) Confocal images in single z-section of immunofluorescence (IF) for RPA70 (upper) and RPA32 (lower) together with TOM20 as well as FISH of mecciND1. Boxed areas are enlarged. Colocalization between RPA70 or RPA32 (G, green), TOM20 (R, red), and mecciND1 (M, magenta) is shown (n=20 randomly selected areas). (D) Representative structured illumination microscopy (N-SIM) image in z-stacks and single z-section of immunofluorescence (IF) for RPA32 together with TOM40 as well as FISH of mecciND1 in fixed HeLa cells. Boxed areas are enlarged. In (C), scale bars, 5 μm and 500 nm (enlarged areas); in (D), scale bars, 2 μm and 200 nm (enlarged areas); in (A) and (B), error bars, s.e.m.; n=3 independent experiments; **P < 0.01; ***P < 0.001, Student's *t*-test.

**Figure S6. mecciND5 interacts with hnRNPA proteins, Related to Figure 2**

(A) The position of antisense oligo and mecciND5 pulldown efficiency are shown; Actin mRNA served as negative control. (B) Homology of hnRNPA1, hnRNPA2B1, and hnRNPA3 proteins and alignment of their RNA recognition motif (RRM). (C) Confocal images in single z-section of immunofluorescence (IF) for hnRNPA1 (upper), hnRNPA2B1 (middle), and hnRNPA3 (lower) together with TOM20 as well as FISH of mecciND5. Boxed areas are enlarged. Colocalization between hnRNPA1, hnRNPA2B1, or hnRNPA3 (G, green), TOM20 (R, red), and mecciND5 (M, magenta) is shown (n=20 randomly selected areas). (D) Representative structured illumination microscopy (N-SIM) image in z-stacks and single z-section of immunofluorescence (IF) for hnRNPA1 together with TOM40 as well as FISH of mecciND5 in fixed HeLa cells. Boxed areas are enlarged. In (C), scale bars, 5 μm and 500nm (enlarged areas). In (D), scale bars, 2 μm and 200 nm (enlarged areas); in (A), error bars, s.e.m.; n=3 independent experiments; ***P < 0.001, Student's *t*-test.

**Figure S7. Correlations between mecciND1 and mitochondrial RPA levels, Related to Figure 3**

(A) Knockdown efficiency of mecciND1 for Figure 3A; si-NC, siRNA with scrambled sequences. (B) Changes in mitochondrial RPA protein levels upon the transfection of antisense morpholino oligos (AMO) against mecciND1 (mecciND1-AMO). Quantification of RPA proteins is shown (normalized to TIM23, a mitochondrial inner membrane protein). Whole cell levels and cytosolic levels of mecciND1 are shown with bar figure. (C) Diagram of mecciND1 overexpression plasmid.

MecciND1 RNA levels increased in both whole cells (total level) and mitochondria (mito level) through the transfected overexpression plasmid (OE-mecciND1). (D) RPA70 and RPA32 mRNA levels were unchanged after the knockdown or overexpression of mecciND1. (E) Knockdown efficiency of mecciND1 for Figure 3C, 3D and Figure S7F; si-NC, siRNA with scrambled sequences. (F) Representative structured illumination microscopy images in z-stacks (3D N-SIM) of immunofluorescence (IF) for PRA32 together with TOM40 as well as FISH of mecciND1 in fixed RPE-1 cells transfected with siRNA (si-NC or si-mecciND1). Single z-section images of boxed areas are enlarged. Scale bars, 2 μm and 200 nm (enlarged areas). In (A)-(E), relative RNA levels were normalized to 18S rRNA (for total level and cytosol level) and 16S rRNA (for mito level), error bars; s.e.m.; n=3 independent experiments; ns, not significant; *P < 0.05; ***P < 0.001, Student's *t*-test.

**Figure S8. Correlations between mecciND5 and mitochondrial hnRNPA levels, Related to Figure 4**

(A) Knockdown efficiency of mecciND5 for Figure 4A, si-NC, siRNA with scrambled sequences; hnRNPA1, hnRNPA2B1, and hnRNPA3 mRNA levels were examined under mecciND5 knockdown. (B) Changes in mitochondrial hnRNPA protein levels upon the transfection of antisense morpholino oligos (AMO) against mecciND5 (mecciND5-AMO). Quantification of hnRNPA proteins is shown (normalized to TIM23, a mitochondrial inner membrane protein). Whole cell levels and cytosolic levels of mecciND5 are shown with bar figure. (C) Diagram of mecciND5 overexpression plasmid. MecciND5 levels increased in both whole cells (total level) and mitochondria (mito level) through plasmid overexpression (OE). (D) hnRNPA1, hnRNPA2B2, and hnRNPA3 mRNA levels upon mecciND5 overexpression. (E) Knockdown efficiency of mecciND5 for Figure 4C, 4D and Figure S8F. (F) Representative structured illumination microscopy images in z-stacks (3D N-SIM) of immunofluorescence (IF) for hnRNPA1 together with TOM40 as well as FISH of mecciND5 in fixed RPE-1 cells transfected with siRNA (si-NC or si-mecciND5). Single z-section images of boxed areas are enlarged. Scale bars, 2 μm and 200 nm (enlarged areas). In (A)-(E), relative RNA levels were normalized to 18S rRNA (total level and cytosol level) and 16S rRNA (mito level); error bars, s.e.m.; n=3 independent experiments; ns, not significant; *P < 0.05; **P < 0.01; ***P < 0.001, Student's *t*-test.

**Figure S9. mecciND1 and mecciND5 copy number per cell and *in vitro* assays, Related to Figure 5**

(A) mecciND1 and mecciND5 copy number per cell in 293T, HeLa, RPE-1, and HepG2 cell lines. (B) Schematic diagram showing procedures of linear RNA generation and *in vitro* circularization[23]. E1, 14 bp of Td gene exon1; E2, 15 bp of Td gene exon2. (C) 5% Urea PAGE gel of purified circular RNAs, mecciND1, mecciND5, and circSRSF. (D) Semi-quantitative RT-PCR of RNA import results for Figure 5B. (E) Semi-quantitative RT-PCR of RNA import results for Figure 5C. (F) Semi-quantitative RT-PCR showed that mecciND1 and mecciND5 imported into mitochondria could be exported out in the *in vitro* assays. Super, supernatant; Pellet, mitochondria.

**Figure S10. Western blots of proteins in the RNA-IP (for Fig. 7a) and knockdown and overexpression of PNPASE, Related to Figure 5**

**(A)** Western blots of overexpressed TOM20-FLAG, TOM40-FLAG and PNPASE-FLAG. NDUFB8, a mitochondrial marker. (B) Western blots to show the successful IP with anti-FLAG antibodies (α-FLAG). Open triangles indicate specific bands of ACTB and * denotes antibody heavy chain. (C) RT-qPCR and Western blots showed the knockdown efficiency of PNPASE for Figure 5G. TOM40, served as loading control. (D) Mitochondrial mecciND1 levels of 293T cells under PNPASE overexpression. RMRP RNA, a known PNPASE interacting RNA imported into mitochondria. Vec, vector control; PNP, PNPASE overexpression. In (C) and (D), relative RNA levels were normalized to 18S rRNA (total level) and 16S rRNA (mito level); error bars, s.e.m.; n=3 independent experiments; ns, not significant; $^{**}P < 0.01$; $^{***}P < 0.001$, Student's t-test.

**Figure S11. Secondary structure of mecciND1 and mecciND5, Related to Figure 5**

(A) A predicted secondary structure of mecciND1 generated by the mfold. Boxed areas are enlarged. (B) *In vivo* secondary structure of mecciND1 sequence in published icSHAPE data (Sun et al., 2019). The underlined sequences corresponded to the boxed stem-loop structure in (A). (C) A predicted secondary structure of mecciND5 generated by the mfold Web Server. Boxed areas are enlarged. (D) *In vivo* secondary structure of mecciND5 sequence in published icSHAPE data. The underlined sequences corresponded to the boxed stem-loop structure in (D).

**Figure S12. mecciND5 levels in HCC, mecciRNAs in *S. pombe* and *C. elegans*, and a working model of mecciRNAs. Related to Figure 6**

(A) mecciND5 levels in pairs of tumor samples and adjacent tissues from 21 Hepatocellular carcinoma (HCC) patients. Relative RNA levels were normalized to 18S rRNA; error bars, s.e.m.; *$P < 0.05$; by Student's *t*-test. (B) Identification of mecciRNAs in *S. pombe* using convergent and divergent primers. g+mtDNA, nuclear and mitochondrial DNA; no RT, reverse transcription reaction without reverse transcriptase. (C) Identification of mecciRNAs in *C. elegans* using convergent and divergent primers. g+mtDNA, nuclear and mitochondrial DNA; no RT, reverse transcription reaction without reverse transcriptase. (D) A working model of mecciRNAs in facilitating mitochondrial protein importation.

### Figure S1

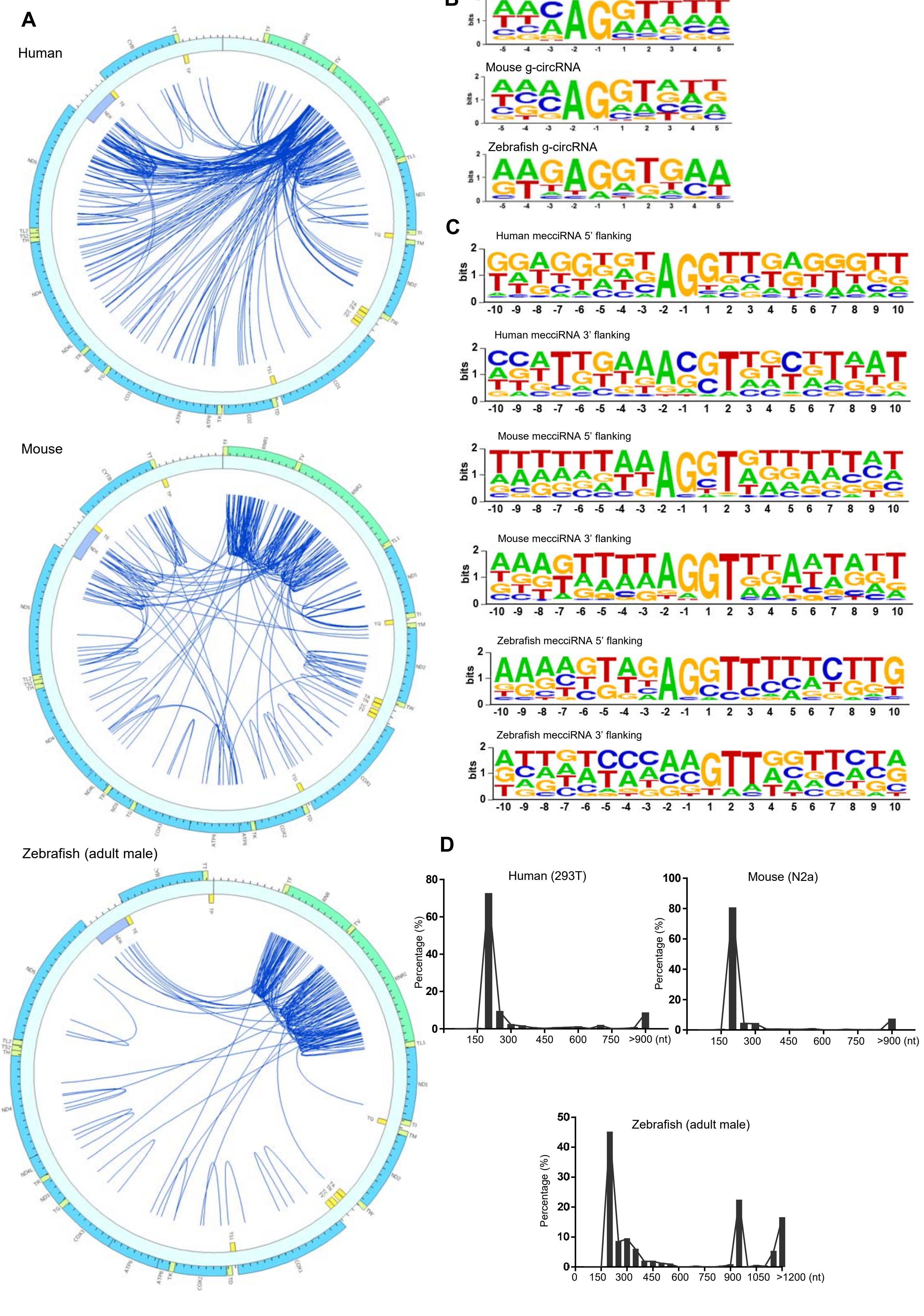
A
Human
Mouse
Zebrafish (adult male)
B
Human g-circRNA
Mouse g-circRNA
Zebrafish g-circRNA
C
Human mecciRNA 5' flanking
Human mecciRNA 3' flanking
Mouse mecciRNA 5' flanking
Mouse mecciRNA 3' flanking
Zebrafish mecciRNA 5' flanking
Zebrafish mecciRNA 3' flanking
D
Human (293T)
Mouse (N2a)
Zebrafish (adult male)
Percentage (%)
150
300
450
600
750
>900 (nt)
1050
>1200 (nt)

### Figure S2

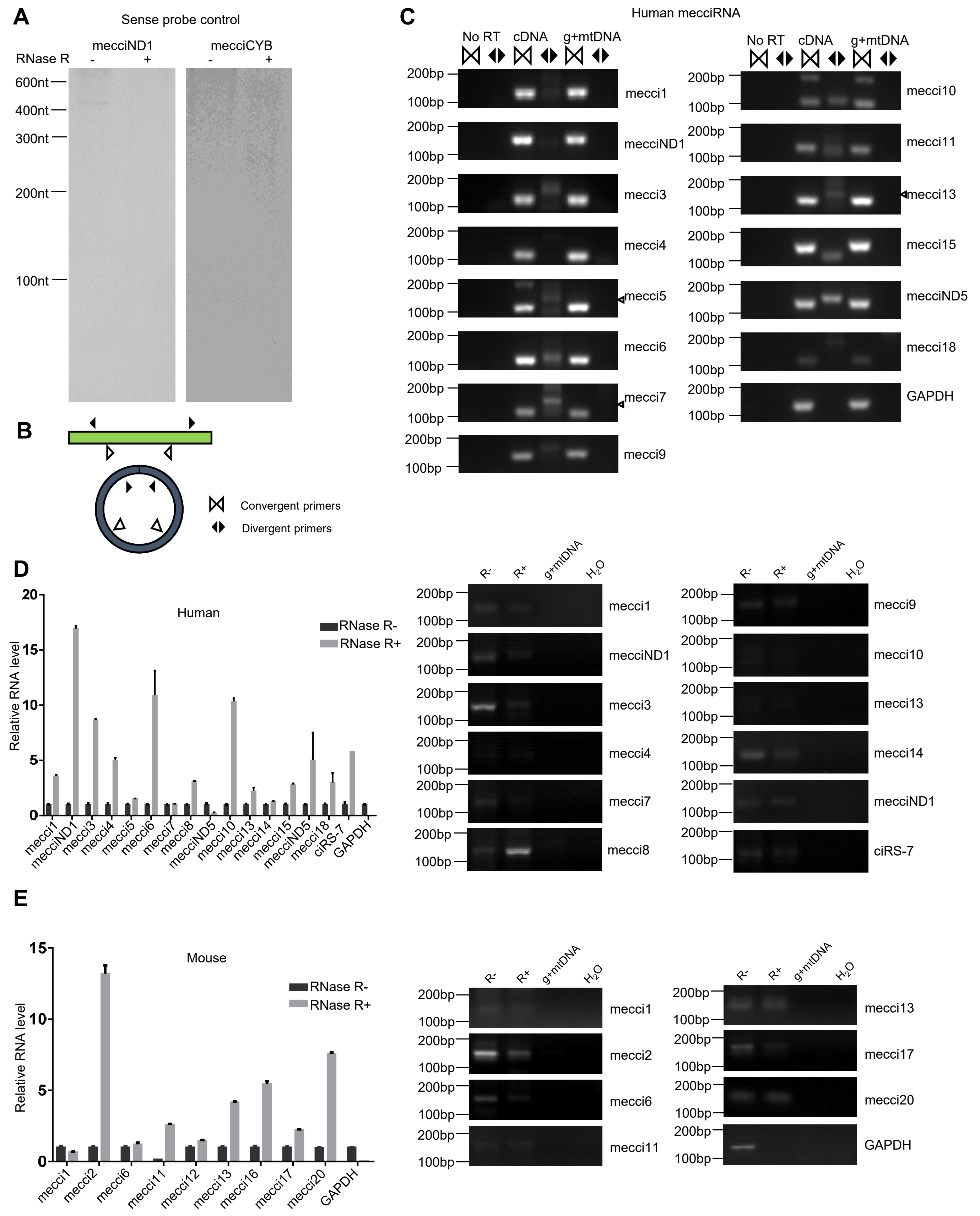

A
Sense probe control
mecciND1
mecciCYB
RNase R
600nt
400nt
300nt
200nt
100nt
B
Convergent primers
Divergent primers
C
Human mecciRNA
No RT
cDNA
g+mtDNA
200bp
100bp
mecci1
mecciND1
mecci3
mecci4
mecci5
mecci6
mecci7
mecci9
mecci10
mecci11
mecci13
mecci15
mecciND5
mecci18
GAPDH
D
Human
RNase R-
RNase R+
Relative RNA level
mecci1
mecciND1
mecci3
mecci4
mecci5
mecci6
mecci7
mecci8
mecciND5
mecci10
mecci13
mecci14
mecci15
mecciND5
mecci18
ciRS-7
GAPDH
R-
R+
g+mtDNA
$H_2O$
mecci9
mecci10
mecci13
mecci14
mecciND1
ciRS-7
E
Mouse
RNase R-
RNase R+
Relative RNA level
mecci1
mecci2
mecci6
mecci11
mecci12
mecci13
mecci16
mecci17
mecci20
GAPDH

### Figure S3

**A**

| Sample | chrM reads | mecciRNA juction reads |
|---|---|---|
| Nuc-DNA | 158025 | 0 |
| Mt-DNA | 34676 | 0 |

**B**

| Sample | Circular RNA analysis | | Linear RNA analysis | | | | | |
|---|---|---|---|---|---|---|---|---|
| | g-circRNA number | mecciRNA number | total reads | mt-mRNA reads | | | | |
| | | | | mt-Cytb | mt-Nd5 | mt-Co3 | mt-Co1 | mt-Nd1 |
| wt MEF (SRR6824985) | 5991 | 9 | 113207764 | 276097 | 203321 | 255401 | 595868 | 334556 |
| Rho0 MEF (SRR6824988) | 6619 | 0 | 111377076 | 0 | 0 | 0 | 0 | 0 |

**C**

| Nascent circRNA identified from Nature Methods, 2018 (HeLa cell) | | |
|---|---|---|
| | circRNA number | circRNA total reads |
| **g-circRNA** | 5794 | 6948 |
| **mecciRNA** | 405 | 1110 |

### Figure S4

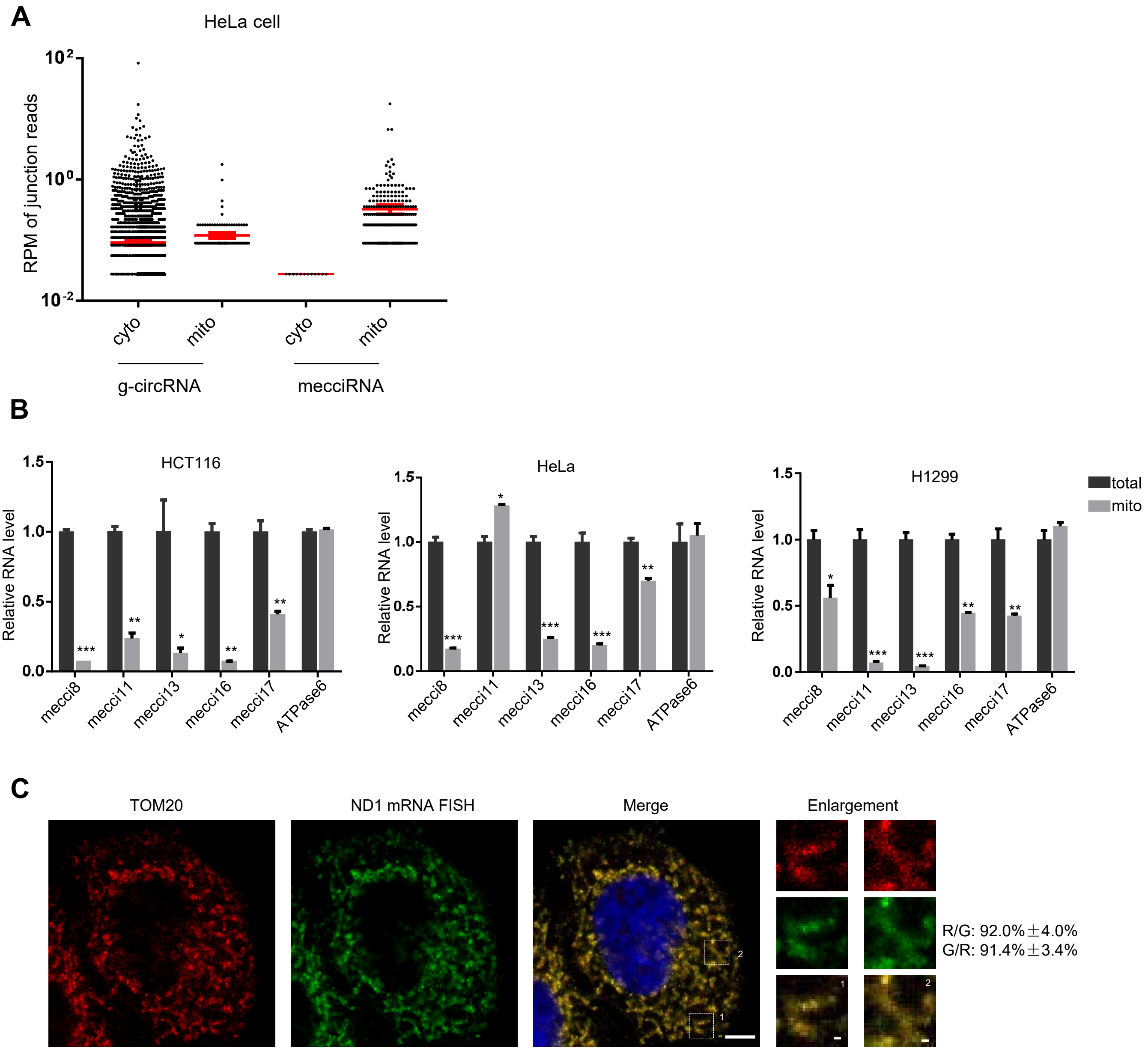
A
HeLa cell
RPM of junction reads
10^2
10^0
10^-2
cyto
mito
cyto
mito
g-circRNA
mecciRNA
B
HCT116
HeLa
H1299
Relative RNA level
1.5
1.0
0.5
0.0
total
mito
mecci8
mecci11
mecci13
mecci16
mecci17
ATPase6
C
TOM20
ND1 mRNA FISH
Merge
Enlargement
R/G: 92.0%±4.0%
G/R: 91.4%±3.4%

### Figure S5

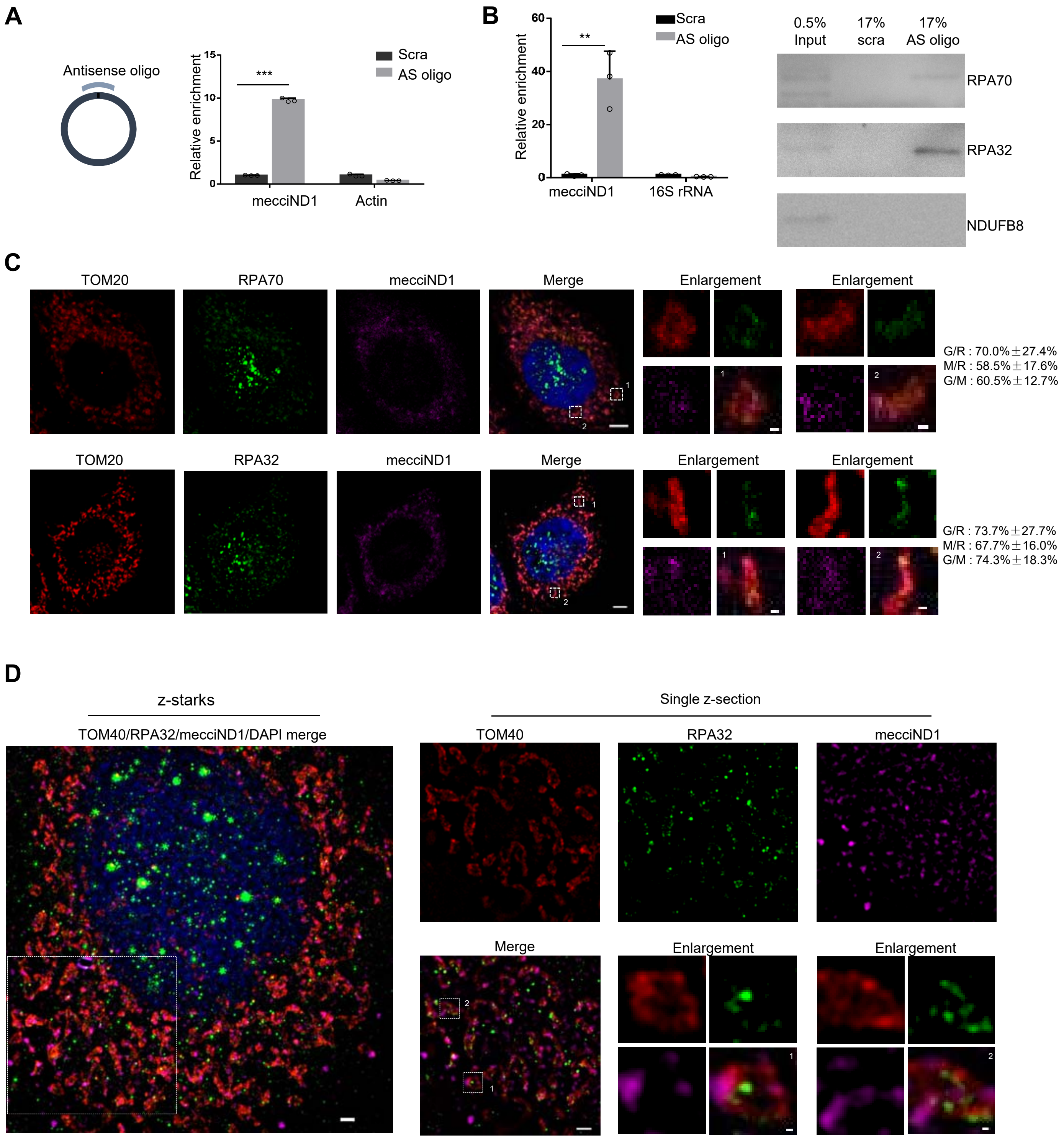

A
Antisense oligo
Relative enrichment
Scra
AS oligo
***
mecciND1
Actin
B
Relative enrichment
Scra
AS oligo
**
mecciND1
16S rRNA
0.5% Input
17% scra
17% AS oligo
RPA70
RPA32
NDUFB8
C
TOM20
RPA70
mecciND1
Merge
Enlargement
G/R : 70.0%±27.4%
M/R : 58.5%±17.6%
G/M : 60.5%±12.7%
TOM20
RPA32
mecciND1
Merge
Enlargement
G/R : 73.7%±27.7%
M/R : 67.7%±16.0%
G/M : 74.3%±18.3%
D
z-starks
TOM40/RPA32/mecciND1/DAPI merge
Single z-section
TOM40
RPA32
mecciND1
Merge
Enlargement

### Figure S6

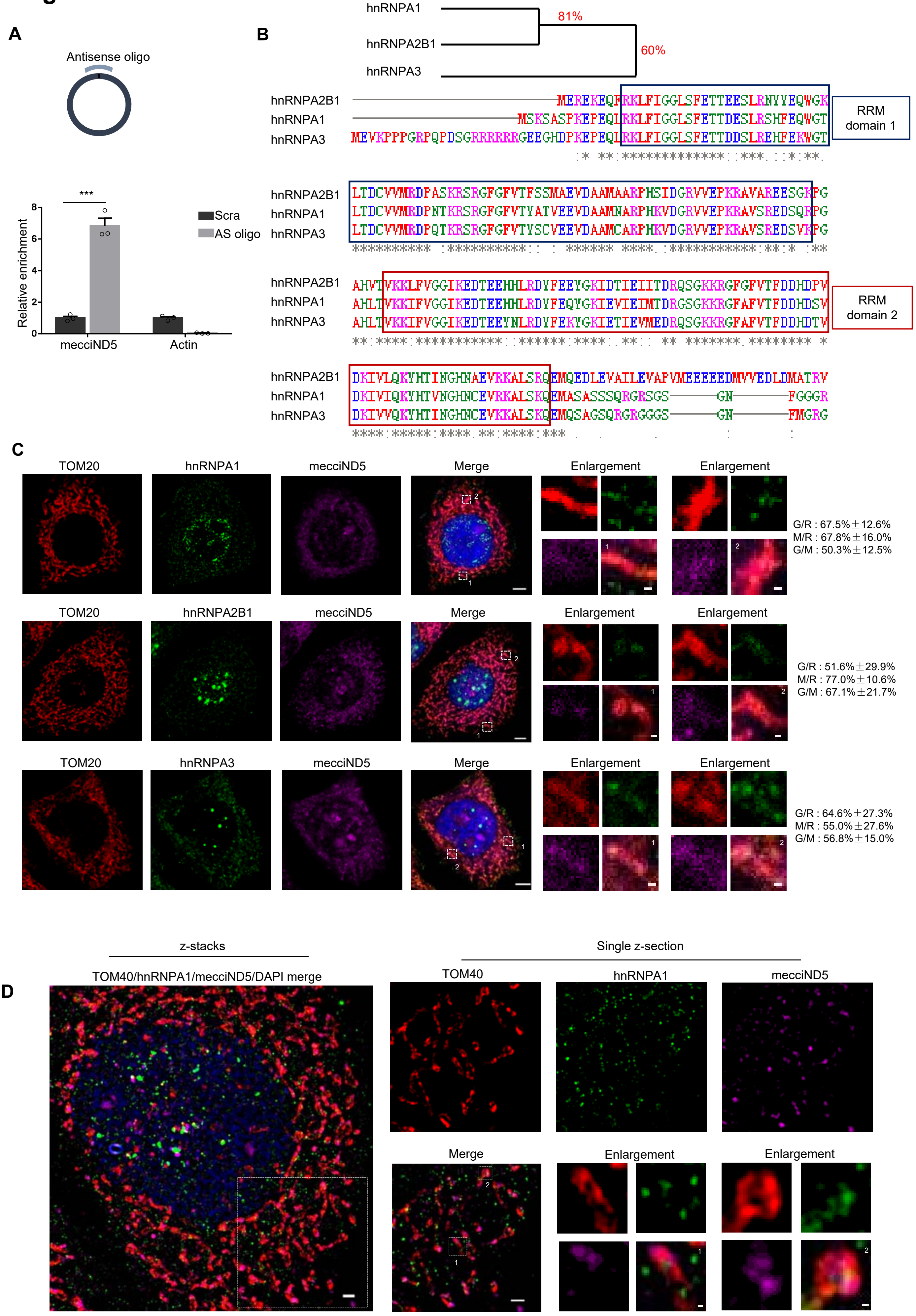

A
Antisense oligo
***
Relative enrichment
8
6
4
2
0
Scra
AS oligo
mecciND5
Actin
B
hnRNPA1
hnRNPA2B1
hnRNPA3
81%
60%
hnRNPA2B1 MEREKEQFRKLFIGGLSFETTEESLRNYYEQWGK
hnRNPA1 MSKSASPKEPEQLRKLFIGGLSFETTDESLRSHFEQWGT
hnRNPA3 MEVKPPPGRPQPDSGRRRRRRGEEGHDPKEPEQLRKLFIGGLSFETTDDSLREHFEKWGT
RRM domain 1
hnRNPA2B1 LTDCVVMRDPASKRSRGFGFVTFSSMAEVDAAMAARPHSIDGRVVEPKRAVAREESGKPG
hnRNPA1 LTDCVVMRDPNTKRSRGFGFVTYATVEEVDAAMNARPHKVDGRVVEPKRAVSREDSQRPG
hnRNPA3 LTDCVVMRDPQTKRSRGFGFVTYSCVEEVDAAMCARPHKVDGRVVEPKRAVSREDSVKPG
hnRNPA2B1 AHVTVKKLFVGGIKEDTEEHHLRDYFEEYGKIDTIEIITDRQSGKKRGFGFVTFDDHDPV
hnRNPA1 AHLTVKKIFVGGIKEDTEEHHLRDYFEQYGKIEVIEIMTDRGSGKKRGFAFVTFDDHDSV
hnRNPA3 AHLTVKKIFVGGIKEDTEEYNLRDYFEKYGKIETIEVMEDRQSGKKRGFAFVTFDDHDTV
RRM domain 2
hnRNPA2B1 DKIVLQKYHTINGHNAEVRKALSRQEMQEDLEVAILEVAPVMEEEEEDMVVEDLDMATRV
hnRNPA1 DKIVIQKYHTVNGHNCEVRKALSKQEMASASSSQRGRSGS GN FGGGR
hnRNPA3 DKIVVQKYHTINGHNCEVKKALSKQEMQSAGSQRGRGGGS GN FMGRG
C
TOM20
hnRNPA1
mecciND5
Merge
Enlargement
G/R : 67.5%±12.6%
M/R : 67.8%±16.0%
G/M : 50.3%±12.5%
hnRNPA2B1
G/R : 51.6%±29.9%
M/R : 77.0%±10.6%
G/M : 67.1%±21.7%
hnRNPA3
G/R : 64.6%±27.3%
M/R : 55.0%±27.6%
G/M : 56.8%±15.0%
D
z-stacks
Single z-section
TOM40/hnRNPA1/mecciND5/DAPI merge
TOM40
hnRNPA1
mecciND5
Merge
Enlargement

### Figure S7

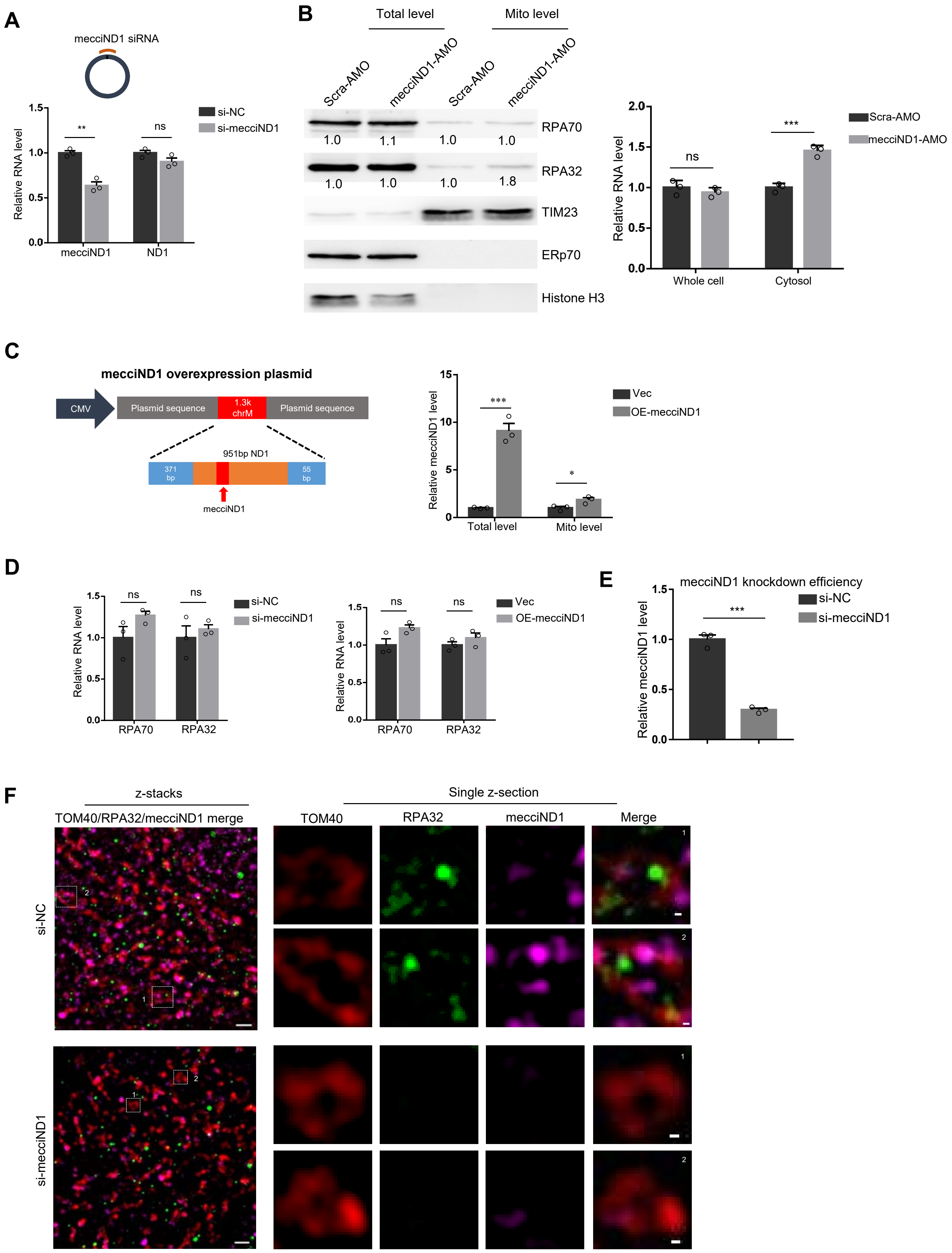
A
mecciND1 siRNA
si-NC
si-mecciND1
Relative RNA level
mecciND1
ND1
**
ns
B
Total level
Mito level
Scra-AMO
mecciND1-AMO
RPA70
RPA32
TIM23
ERp70
Histone H3
1.0
1.1
1.8
Whole cell
Cytosol
***
C
mecciND1 overexpression plasmid
CMV
Plasmid sequence
1.3k chrM
951bp ND1
371 bp
55 bp
mecciND1
Vec
OE-mecciND1
Relative mecciND1 level
Total level
Mito level
*
D
RPA70
RPA32
E
mecciND1 knockdown efficiency
F
z-stacks
TOM40/RPA32/mecciND1 merge
Single z-section
TOM40
RPA32
mecciND1
Merge
si-NC
si-mecciND1

### Figure S8

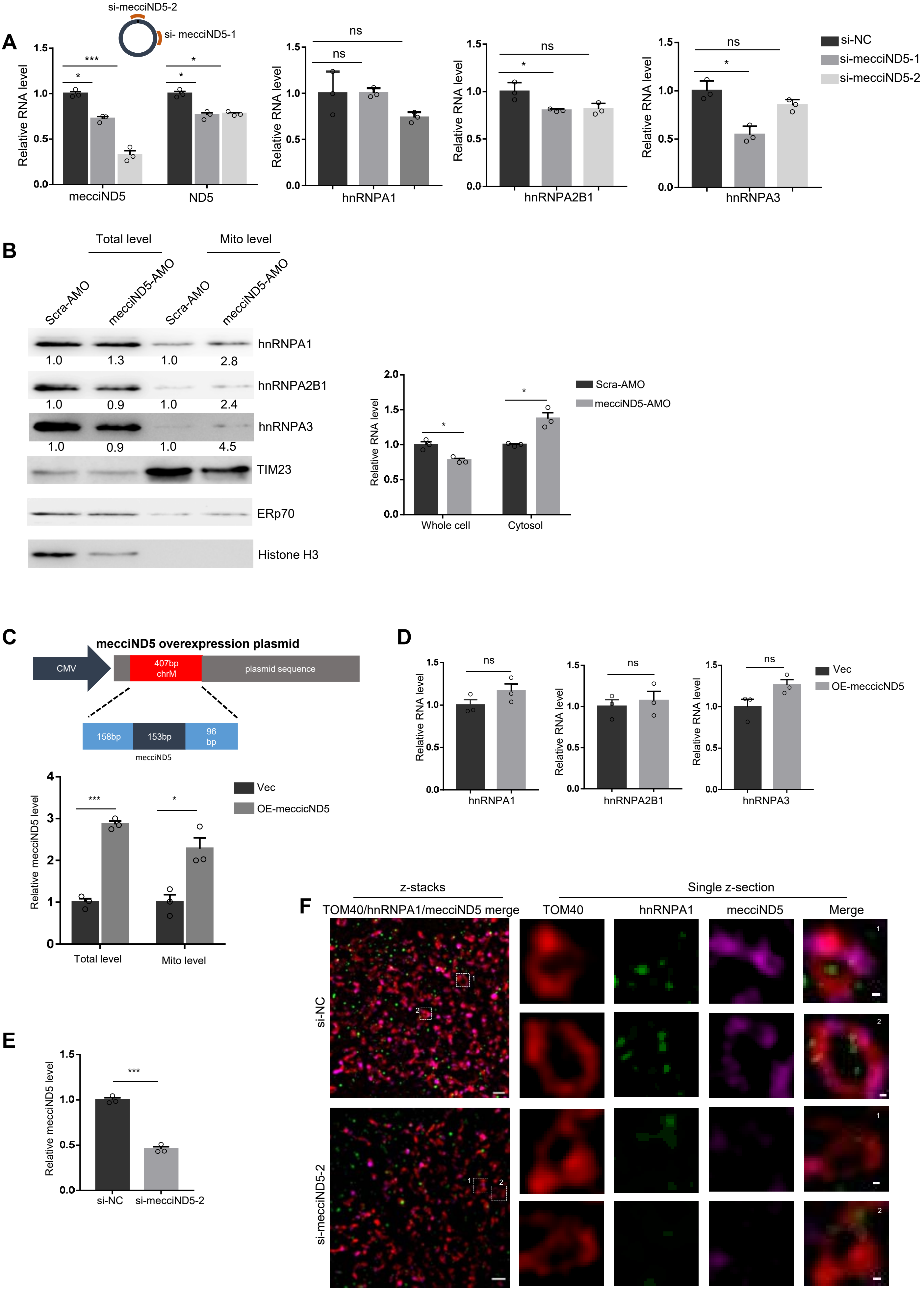
A
si-mecciND5-2
si- mecciND5-1
Relative RNA level
mecciND5
ND5
hnRNPA1
hnRNPA2B1
hnRNPA3
si-NC
si-mecciND5-1
si-mecciND5-2
ns
B
Total level
Mito level
Scra-AMO
mecciND5-AMO
hnRNPA1
hnRNPA2B1
hnRNPA3
TIM23
ERp70
Histone H3
1.0 1.3 1.0 2.8
1.0 0.9 1.0 2.4
1.0 0.9 1.0 4.5
Whole cell
Cytosol
C
mecciND5 overexpression plasmid
CMV
407bp chrM
plasmid sequence
158bp
153bp
96 bp
mecciND5
Relative mecciND5 level
Vec
OE-meccicND5
Total level
Mito level
D
E
si-NC
si-mecciND5-2
F
z-stacks
Single z-section
TOM40/hnRNPA1/mecciND5 merge
TOM40
hnRNPA1
mecciND5
Merge
si-NC
si-mecciND5-2

### Figure S9

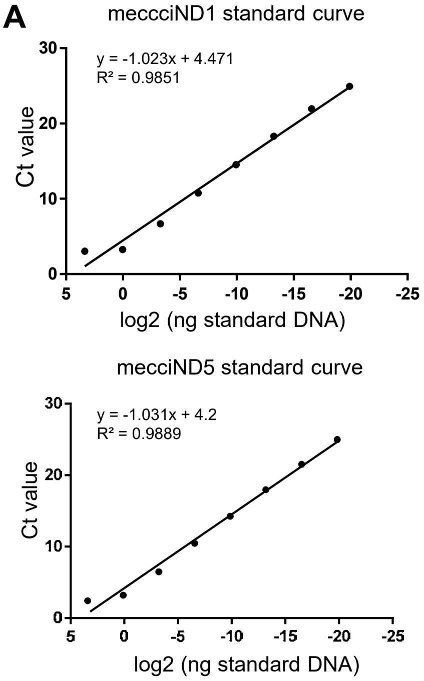

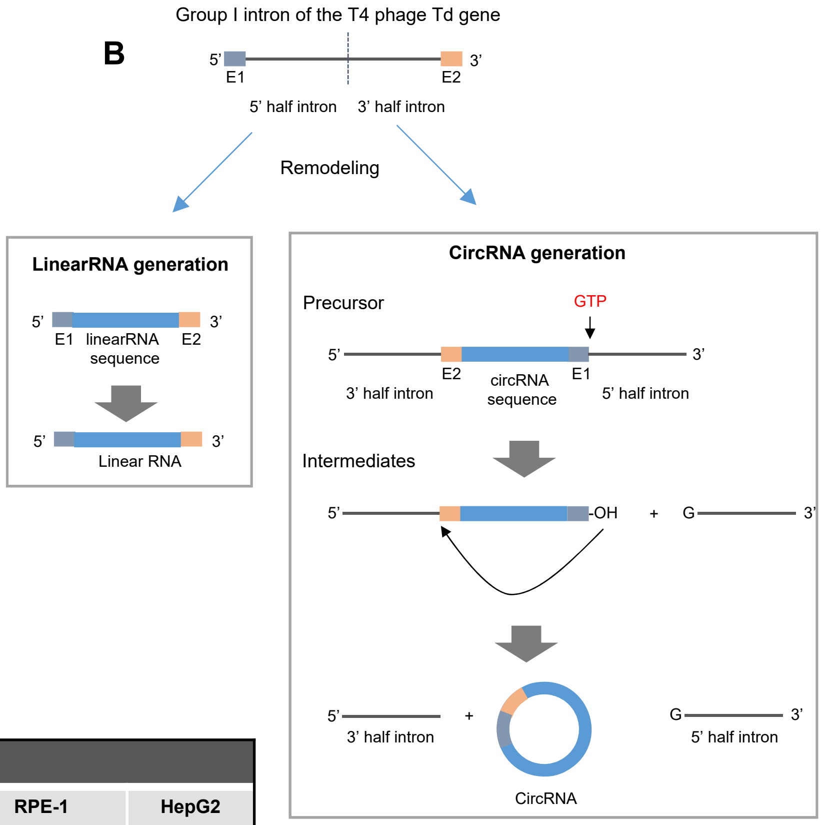

| RNA copy number per cell | | | | |
|---|---|---|---|---|
| | **293T** | **HeLa** | **RPE-1** | **HepG2** |
| **mecciND1** | ~74 | ~61 | ~52 | ~109 |
| **mecciND5** | ~195 | ~330 | ~118 | ~222 |

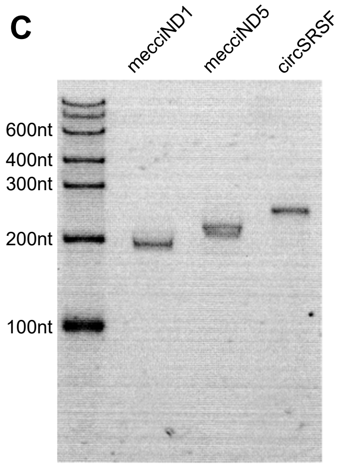

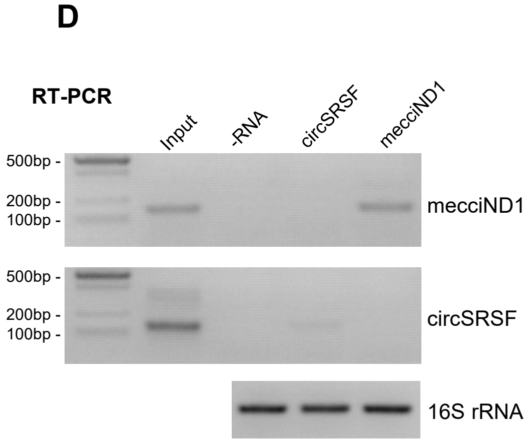

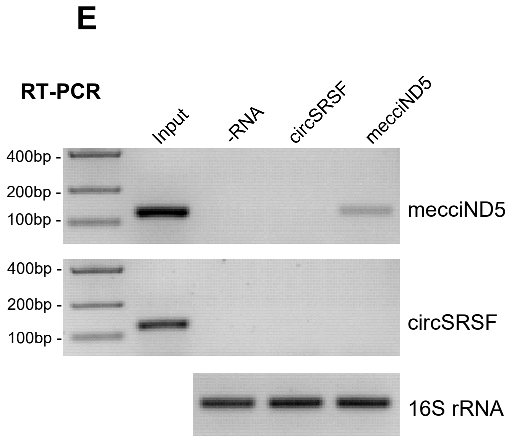

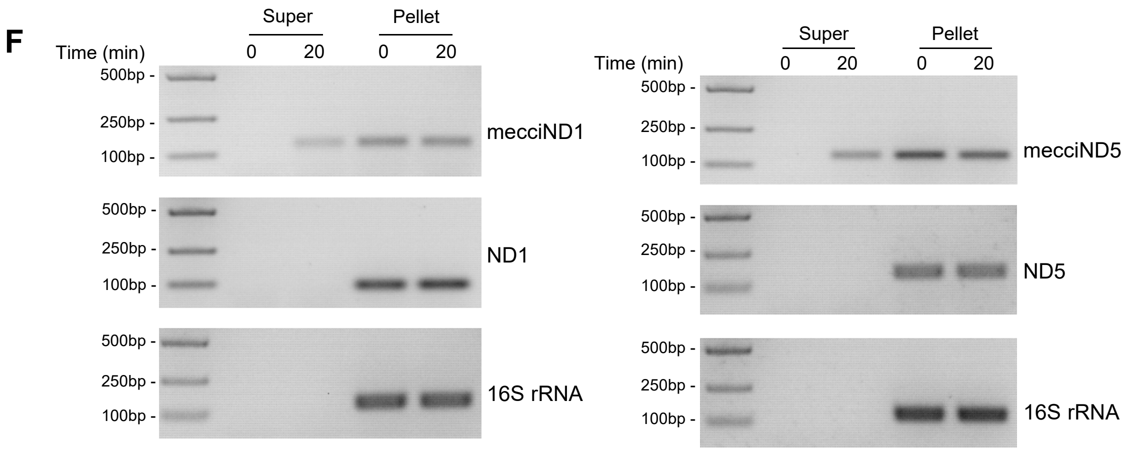

### Figure S10

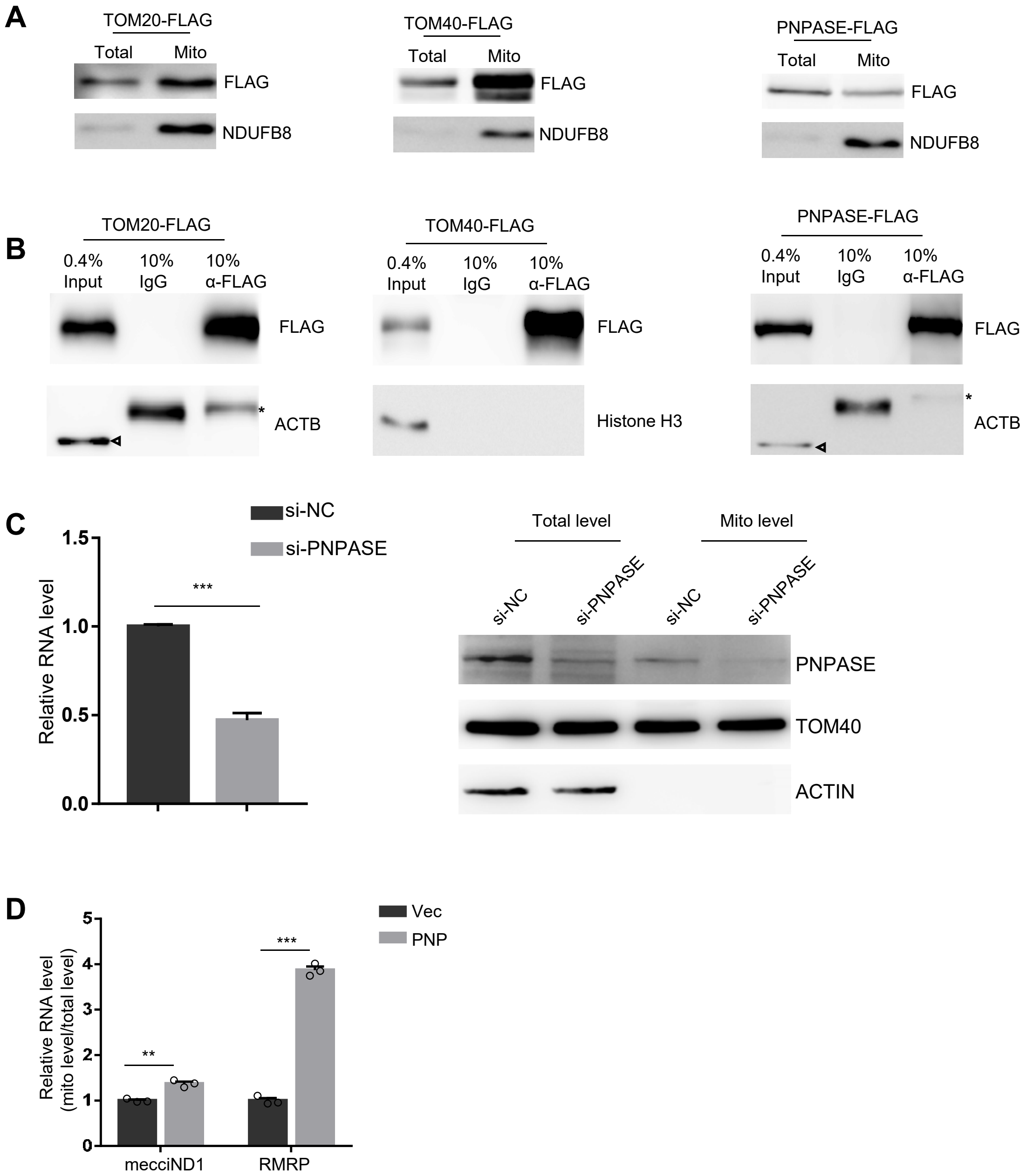

A
TOM20-FLAG
Total
Mito
FLAG
NDUFB8
TOM40-FLAG
Total
Mito
FLAG
NDUFB8
PNPASE-FLAG
Total
Mito
FLAG
NDUFB8
B
TOM20-FLAG
0.4% Input
10% IgG
10% α-FLAG
FLAG
ACTB
TOM40-FLAG
0.4% Input
10% IgG
10% α-FLAG
FLAG
Histone H3
PNPASE-FLAG
0.4% Input
10% IgG
10% α-FLAG
FLAG
ACTB
C
si-NC
si-PNPASE
Relative RNA level
1.5
1.0
0.5
0.0
***
Total level
Mito level
si-NC
si-PNPASE
si-NC
si-PNPASE
PNPASE
TOM40
ACTIN
D
Vec
PNP
Relative RNA level (mito level/total level)
5
4
3
2
1
0
**
***
mecciND1
RMRP

### Figure S11

A

mecciND1

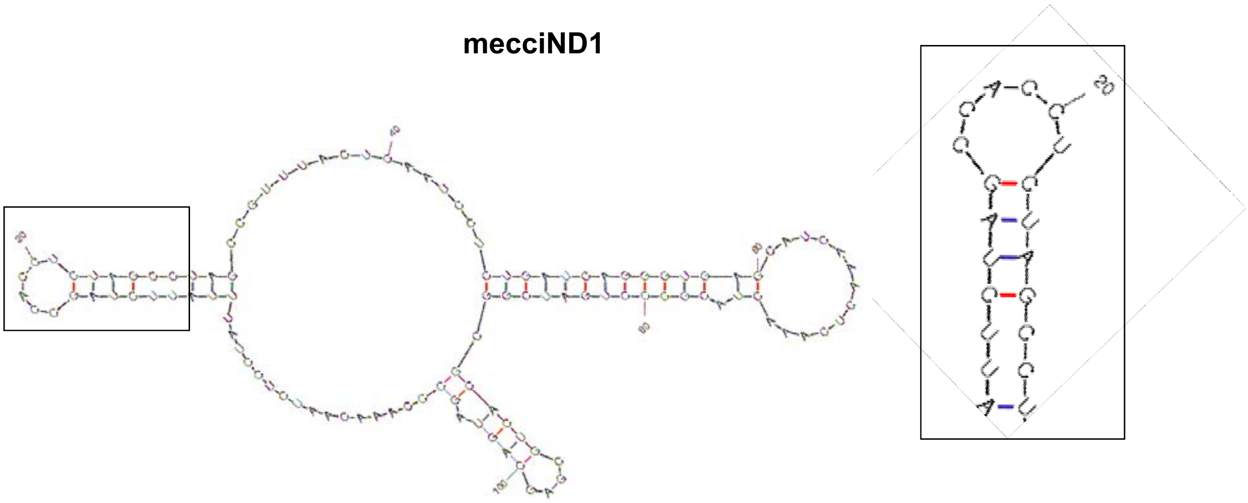

B

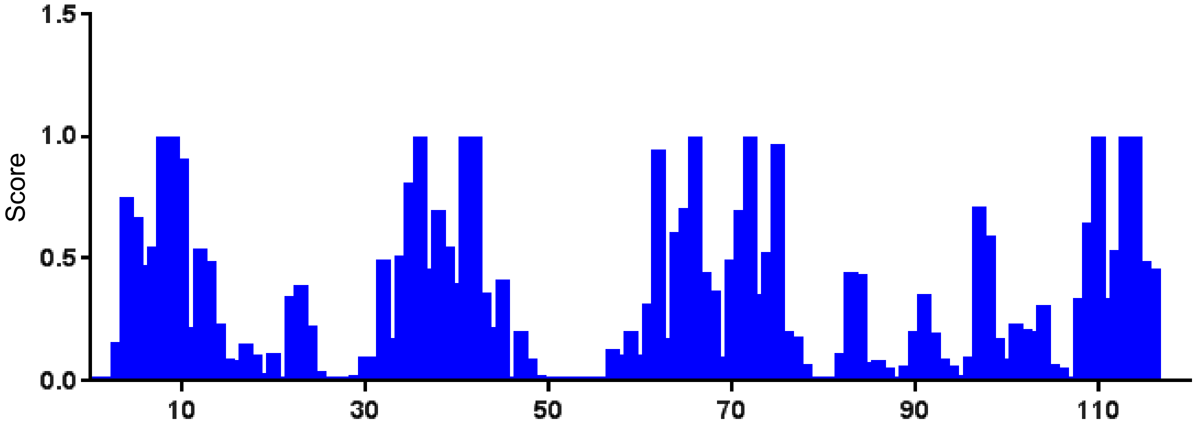

TCCTATTTATTCTAGCCACCTCTAGCCTAGCCGTTTACTCAATCCTCTGATCAGGGTGAGCATCAAACTCAAACTACGCCCTGATCGGCGCACTGCGAGCAGTAGCCCAAACAATC

C

mecciND5

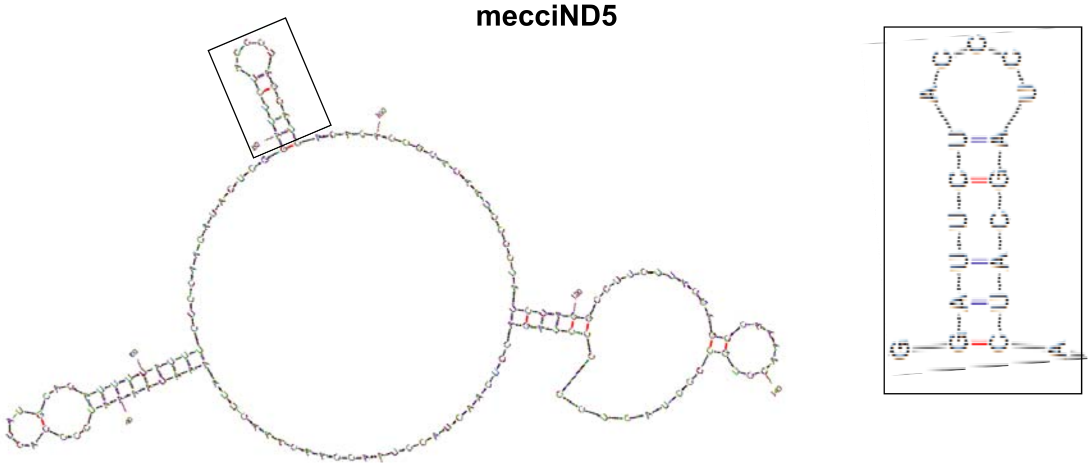

D

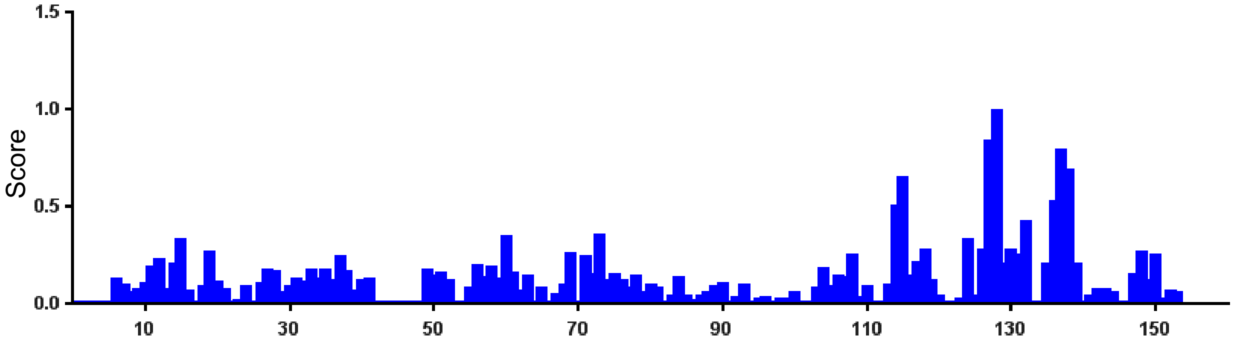

CCTCAACTACCTAACCAACAAACTTAAAATAAAATCCCCACTATGCACATTTTATTTCTCCAACATACTCGGATTCTACCCTAGCATCACACACCGCACAATCCCCTATCTAGGCCTTCTTACGAGCCAAAACCTGCCCCTACTCCTCCTAGAC

### Figure S12

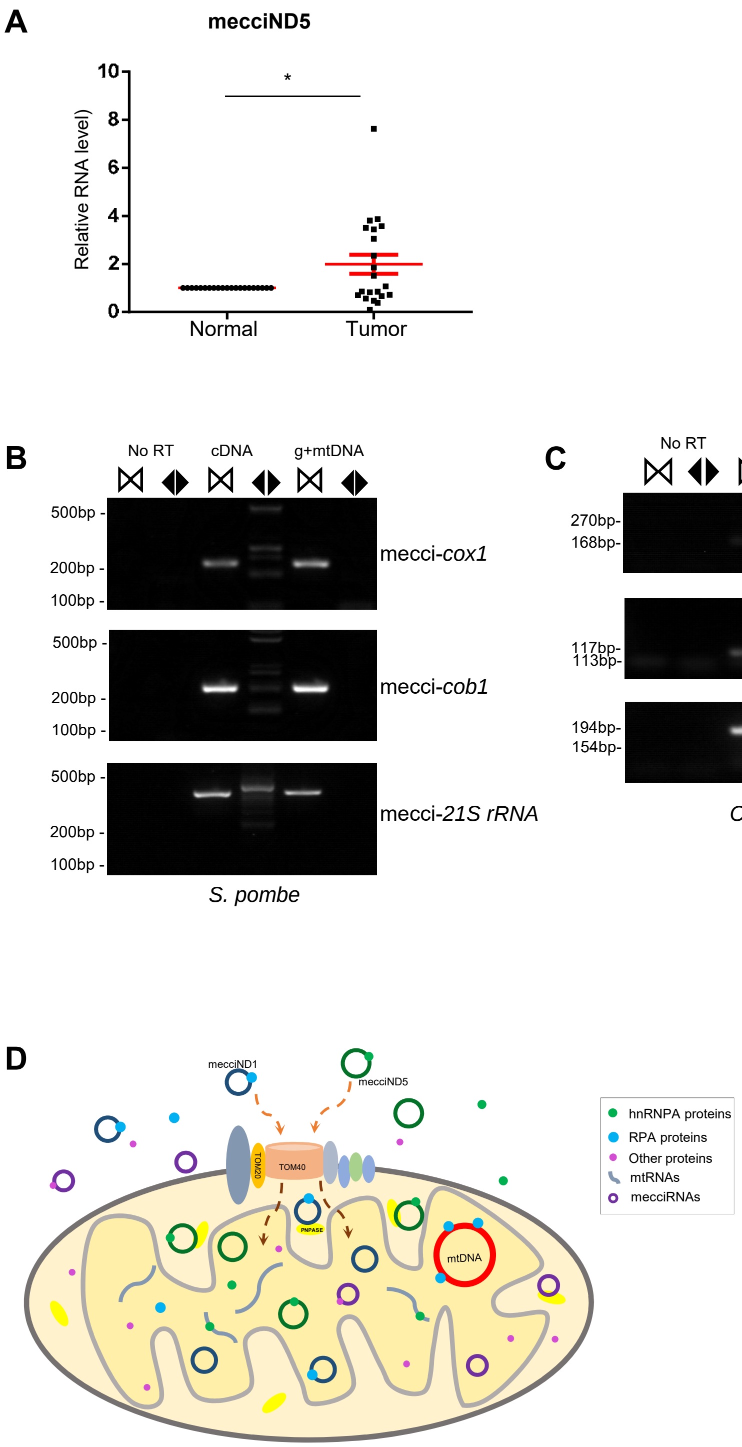

A
mecciND5
Relative RNA level)
10
8
6
4
2
0
*
Normal
Tumor
B
No RT
cDNA
g+mtDNA
500bp -
200bp -
100bp -
mecci-cox1
mecci-cob1
mecci-21S rRNA
S. pombe
C
270bp-
168bp-
117bp-
113bp-
194bp-
154bp-
mecci-nduo-1
mecci-nduo-5
mecci-ctc-1
C. elegans
D
mecciND1
mecciND5
TOM20
TOM40
PNPASE
mtDNA
hnRNPA proteins
RPA proteins
Other proteins
mtRNAs
mecciRNAs

| Table S1 Oligos used in this study | | | |
|---|---|---|---|
| **human mecciRNA detection primer** | | | |
| name | forward primer 5'-3' | reverse primer 5'-3' | |
| h_con_mecci1 | CCAACCCCTTAAACACCCCT | TAGTAATAGGGCAAGGACGC | for human mecciRNA PCR and realtime-PCR |
| h_div_mecci1 | CCGATCCGTCCCTAACAAAC | GAATTGTGTAGGCGAATAGG | |
| h_con_mecci3 | CCTAACCCTGACTTCCCTAA | AGGTGGATGCGACAATGGAT | |
| h_div_mecci3 | CCATTGTCGCATCCACCT | GTTAACGAGGGTGGTAAGGA | |
| h_con_mecci4 | CAAGTATTGACTCACCCATC | GGTGGTCAAGTATTTATGGT | |
| h_div_mecci4 | TACTGCCAGCCACCATGAAT | GAAATACATAGCGGTTGTTG | |
| h_con_mecci5 | CTATTCGCCTACACAATTCTC | AAAGTGATTGGCTTAGTGG | |
| h_div_mecci5 | CCACTAAGCCAATCACTTT | GAGAATTGTGTAGGCGAATAG | |
| h_con_mecci6 | CTAGCCACCTCTAGCCTAGC | GTTTGGGCTACTGCTCGCAG | |
| h_div_mecci6 | GCATCAAACTCAAACTAC | ATCAGAGGATTGAGTAA | |
| h_con_mecci8 | TCCATTGTCGCATCCACCTT | TGGCTCAGTGTCAGTTCGA | |
| h_div_mecci8 | TCGAACTGACACTGAGCCA | AAGGTGGATGCGACAATGGA | |
| h_con_mecci9 | AACAACAACCTATTTAGC | CGTGATAGTGGTTCGCTGG | |
| h_div_mecci9 | CAGGCACATACTTCCTATTC | GCTAAATAGGTTGTTGTTG | |
| h_con_mecci10 | ATAACCCAATACCAAACGCC | TTAGTAGTATAGTGATGCC | |
| h_div_mecci10 | CAGTCCTAGCTGCTGGCATC | AGTAGGACTGCTGTGATTAG | |
| h_con_mecci11 | CTCCAACATACTCGGATTCT | GATTTGGTGCTGTGAAATTG | |
| h_div_mecci11 | CTACTCCTCCTAGACCTAAC | TGTGCGGTGTGTGATGCTAG | |
| h_con_mecci13 | CACCTACTCATGCACCTAAT | GACAGCGATTTCTAGGATAG | |
| h_div_mecci13 | CTATCCTAGAAATCGCTGTC | GAAGATGATAAGTGTAGAGG | |
| h_con_mecci14 | CCTATACTCCCTCTACATAT | AGGAGAATGGGGGATAGGTG | |
| h_div_mecci14 | AACCCTCATTCACACGAGA | TTAATGTGGTGGGTGAGTGAG | |
| h_con_mecci15 | CATGTGCCTAGACCAAGAAG | CTATGATGGACCATGTAACG | |
| h_div_mecci15 | CGTTACATGGTCCATCATAG | GCTTCTTGGTCTAGGCACAT | |
| h_con_mecci18 | ACTCCACCTCAATCACACTA | TAGGTAGGAGTAGCGTGGT | |
| h_div_mecci18 | ACCACGCTACTCCTACCTA | TAGTGTGATTGAGGTGGAGT | |
| h_con_mecciND1 | ACCTCAACCTAGGCCTCCTA | CATATGAGATTGTTTGGGCT | |
| h_con_mecciND1 | TGAGCATCAAACTCAAACTAC | CTAGGCTAGAGGTGGCTAGAA | |
| h_con_mecciND5 | CTCAACTACCTAACCAACAA | TAAGAAGGCCTAGATAGGGG | |
| h_div_mecciND5 | CATCACACACCGCACAATC | AGAATCCGAGTATGTTGGAG | |
| h_ciRS-7 | AACTACCCAGTCTTCCATCA | AGACTTGAAGTCGCTGGAAG | |
| **mouse mecciRNA detection primer** | | | |
| name | forward primer 5'-3' | reverse primer 5'-3' | |
| m_div_mecci1 | GCTTAAGACACCTTGCCTA | TACACCGGTCTATGGAGGTT | for mouse mecciRNA PCR and realtime-PCR |
| m_div_mecci2 | TTTAGATTATAGCCAAAAGAGGGACA | TTTTTGGGTAACCAGCTATCAC | |
| m_div_mecci6 | AGCTAGAAACCCCGAAACCA | TTCATTATGCAAAAGGTACAAGG | |
| m_div_mecci11 | GTGGGCAATTGATGAATAGGC | TCTTCCTTACAACCCATCCCT | |
| m_div_mecci12 | CAGGCAGTGCCTCTAATACT | CATGAACGGCTAAACGAGGG | |
| m_div_mecci13 | GCCACATAGACGAGTTGATTC | AGAGGGACAGCTCTTCTGGAA | |
| m_div_mecci16 | GGTAACTTGGTCCGTTGATC | GGGATAACAGCGCAATCCTA | |
| m_div_mecci17 | TATCCTGACCGTGCAAAGGT | CAGGCAGTGCCTCTAATACT | |
| m_div_mecci20 | GGATTGCGCTGTTATCCCTA | CAGGACATCCCAATGGTGTAG | |
| **real-time qPCR primer** | | | |
| Name | forward primer 5'-3' | reverse primer 5'-3' | |
| q_h/m GAPDH | CTTCATTGACCTCAACTACATGG | CTCGCTCCTGGAAGATGGTGAT | used for both human and mouse |
| q_18S rRNA | CGGCGACGACCCATTCGAAC | GAATCGAACCCTGATTCCCCGTC | |
| q_Actin | GAGTACTTGCGCTCAGGAG | CCAACACAGTGCTGTCTGG | |
| h_q_ND1 | CCCTAAAACCCGCCACATCTA | GAGCGATGGTGAGAGCTAAGGT | |
| h_q_mecciND5 | ATCTAGGCCTTCTTACGAGC | ATTGTGCGGTGTGTGATGCT | |
| h_q_ND5 | GCAGCCATTCAAGCAATCCTA | AGGCGAGGATGAAACCGATA | |
| h_q_ATP6 | TCGGTTGTTGATGAGATATTTGGA | CGCCGCAGTACTGATCATTCT | |
| h_q_12s rRNA | TAGAGGAGCCTGTTCTGTAATCGAT | CGACCCTTAAGTTTCATAAGGGCTA | |
| h_q_RPA70 | GGGGATACAAACATAAAGCCCA | CGATAACGCGGCGGACTATT | |
| h_q_RPA32 | GGTAGCCTTTAAGATCATGCCC | CTGTTGGCTTTGCTTAGTACCA | |
| qh_mtDNA_UUR | CACCCAAGAACAGGGTTTGT | TGGCCATGGGTATGTTGTTA | used for mtDNA copy number |
| qh_nucDNA_B2M | TGCTGTCTCCATGTTTGATGTATCT | TCTCTGCTCCCCACCTCTAAGT | |
| h_q_hnRNPA1 | CCACGAAACCAAGGTGGCTA | TCCCTGTCACTTCTCTGGCT | |
| h_q_hnRNPA2B1 | AGAGGCTTTGGCTTTGTTAC | CCACTCCTAGAACTCTGAAC | |
| h_q_hnRNPA3 | TGGAAGAAGCTCGGGCAGT | ACCTGCAGCTTTCCTGACAA | |
| h_q_PNPASE | CTGCACTACGAGTTTCCTCC | GACCCATTTGACTCTAGGAC | |
| h_q_RMRP | CAGAGAGTGCCACGTGCATA | CTAGAGGGAGCTGACGGATG | used for *in vitro* assay |
| h_16S | ACCAGACGAGCTACCTAAGA | CTTGGACAACCAGCTATCAC | |
| E1-linear ND1-E2 | CTACCGTTTAATATTCTCCT | ACCCAAGAAAACATATTGTGG | |
| E1-linear ND5-E2 | CTACCGTTTAATATTCCCTAG | CCCAAGAAAACATGGAGTAG | |
| div_E2-mecciND1-E1 | CTAGCCTAGCCGTTTACTC | GGAGAATATTAAACGGTAGACC | |
| div_E2-mecciND5-E1 | CTCCAACATACTCGGATTCT | CTAGGGAATATTAAACGGTAGAC | |
| div_E2-circSRSF-E1 | GGATGGAACTGAAGTCAATG | AATCAATATTAAACGGTAGACCC | |
| ***C. elegans* mecciRNA PCR primer** | | | |
| Name | forward primer 5'-3' | reverse primer 5'-3' | |

| | | | |
|---|---|---|---|
| con_mecci-*ctc-1* | TCATAAAGATATCGGAACTC | CGATTATAGTAGGTATTACC | used for *C.elegans* mecciRNA identification |
| div_mecci-*ctc-1* | CGTTTAGAATTAGCTAAACC | CCAACCATACCAGATCAAAG | |
| con_mecci-*nudo-5* | GGCCTATTTACTATATTTTT | CAAAGTAACTATTGAAAAAC | |
| div_mecci-*nduo-5* | CAATAGTTACTTTGGGCCTA | CTATTTGTGAAAGTGTCCTC | |
| con_mecci-*nudo-1* | GGGCCCACCAAGGTTACA | GGTATAATTGGGGCCATC | |
| div_mecci-*nduo-1* | ACTTGTACCAGGAATTTC | CCCATCCAATAAAGCTTG | |
| **S. pombe mecciRNA PCR primer** | | | |
| Name | forward primer 5'-3' | reverse primer 5'-3' | |
| con-pombe-mecci-*cox1* | AATAGGCCTCTTAACGTTGCT | TTGTTGTGATTTCGTTGCGTA | used for *S. pombe* mecciRNA identification |
| div-pombe-mecci-*cox1* | CCTCAGAGACTTTACGCAACG | CAAGTAGTTCCAGCATATAGC | |
| con-pombe-mecci-*21SrRNA* | GTTCAGTATAGAGGTTAGTCG | TAACTTCGCTCATTGAGCACA | |
| div-pombe-mecci-21*SrRNA* | CTCTGTTTGACACCTCGATG | CGACTAACCTCTATACTGAAC | |
| con-pombe-mecci-*cob1* | GAGCTGTATTGCCCGAATTCC | GACCGCATAGTCATTACTGACC | |
| div-pombe-mecci-*cob1* | AGGATACACACCAGGAGAAG | CAGCTAAGACAATCACCTATC | |
| **primers for plasmids** | | | |
| mecciND1-OE-F | ATTCTGCAGTCGACGGTACCGGATCAG GACATCCCGATGG | | mecciND1 overexpression plasmid |
| mecciND1-OE-R | TATCTAGATCCGGTGGATCCGTTTAAGC TCCTATTATTTA | | |
| mecciND5-OE-F | GATCCGCTAGCGCTACCGGTACCCTACT AAACCCCATTA | | mecciND5 overexpression plasmid |
| mecciND5-OE-R | TTATCTAGATCCGGTGGATCCTTATGCC TTTTTGGGTTGAG | | |
| BglII-EcoRI-3xflag-F | GATCTATGGACTACAAAGACCATGACG GTGATTATAAAGATCATGACATCGATTA CAAGGATGACGATGACAAGTAAG | | C-terminal FLAG-tagged human TOM20, TOM40, PNPASE plasmid construction |
| BglII- EcoRI -3xflag-R | AATTCTTACTTGTCATCGTCATCCTTGT AATCGATGTCATGATCTTTATAATCACC GTCATGGTCTTTGTAGTCCATA | | |
| AgeI-TOM20-F | GTCAGATCCGCTAGCGCTACCGGTATG GTGGGTCGGAACAGCGC | | |
| TOM20-BglII-R | GAATTCGAAGCTTGAGCTCGAGATCTTT CCACATCATCTTCAGCCA | | |
| AgeI-TOM40-F | GTCAGATCCGCTAGCGCTACCGGTATG GGGAACGTGTTGGCTGC | | |
| TOM40-BglII-R | GAATTCGAAGCTTGAGCTCGAGATCTG CCGATGGTGAGGCCAAAGC | | |
| BglII-BamHI-3xflag-F | GATCTATGGACTACAAAGACCATGACG GTGATTATAAAGATCATGACATCGATTA CAAGGATGACGATGACAAGTAAG | | |
| BglII- BamHI -3xflag-R | GATCCTTACTTGTCATCGTCATCCTTGT AATCGATGTCATGATCTTTATAATCACC GTCATGGTCTTTGTAGTCCATA | | |
| AgeI-PNPASE-F | GTCAGATCCGCTAGCGCTACCGGTATG GCGGCCTGCAGGTACTG | | |
| PNPASE-BglII-R | CGAAGCTTGAGCTCGAGATCTCTGAGA ATTAGATGATGAC | | |
| Group I intron sequence of Td g | GGTTCTACATAAATGCCTAACGACTATCCCTTTGGGGAGTAGGGTCAAGTGAC TCGAAACGATAGACAACTTGCTTTAACAAGTTGGAGATATAGTCTGCTCTGCA TGGTGACATGCAGCTGGATATAATTCCGGGGTAAGATTAACGACCTTATCTGA ACATAATGCTACCGTTTAATATTATGTTTTCTTGGGTTAATTGAGGCCTGAGTA TAAGGTGACTTATACTTGTAATCTATCTAAACGGGGAACCTCTCTAGTAGACA ATCCCGTGCTAAATTGTAGGACT | | for *in vitro* transcription by T7 promoter and circularization |
| C-T7-E1-F | TGTAATACGACTCACTATAGGTTCTACA TAAATGCCTAA | | |
| C-E2-R | AGTCCTACAATTTAGCACGG | | |
| L-T7-E1-F | TGTAATACGACTCACTATACTACCGTTT AATATT | | |
| L-E2-R | ACCCAAGAAAACAT | | |
| SP6_EcoRI_FLAG_F | GCCGCCAGTGTGCTGGAATTC TTActtgtcatcgtcatc | | plamids for in vitro transcription by SP6 promoter and in vitro translation |
| SP6_RPA2_XbalI_R | TATAGAATAGGGCCCTCTAGAGCCACC ATGTGGAACAGT | | |
| SP6_ HNRNPA1_XbalI_R | TATAGAATAGGGCCCTCTAGAGCCACC ATGTCTAAGTCA | | |
| **oligos and probes** | | | |
| Bio-scramble-oligo | TTCTCCGAACGTGTCACGTTCGAACGTG TC | | probe for Biotin oligo pull down |
| Bio-mecciND1-oligo | CTAGAATAAATAGGAGATTGTTTGG GCTAC | | |
| biotin_mecciND5_probe | AGGTAGTTGAGGTCTAGGGGGAGTAGGG GC | | |

| | | | |
|---|---|---|---|
| mecciND1-s | TGTAATACGACTCACTATAGGGTGAGCATCAAACTCAAACTAC | CTAGGCTAGAGGTGGCTAGAA | primer for Northern blot or FISH probe |
| mecciND1-as | TGAGCATCAAACTCAAACTAC | TGTAATACGACTCACTATAGGGCTAGGCTAGAGGTGGCTAGAA | |
| ND1_out_as | CTCGTTGTACCCATTCTAATCG | TGTAATACGACTCACTATAGGGGTCAGCGAAGGGTTGTAGTAGC | |
| mecciND5_s | TGTAATACGACTCACTATAGGGCCGCACAATCCCCTATCTAG | TGTGTGATGCTAGGGTAGAA | |
| mecciND5_as | CCGCACAATCCCCTATCTAG | TGTAATACGACTCACTATAGGGTGTGTGATGCTAGGGTAGAA | |
| mecci-CYB | CCGATCCGTCCCTAACAAAC | GAATTGTGTAGGCGAATAGG | |
| si-NC | UUCUCCGAACGUGUCACGU | ACGUGACACGUUCGGAGAA | siRNA for mecciRNA knockdown |
| mecciND1-siRNA | CCAAACAAUCUCCUAUUUAU | AUAAAUAGGAGAUUGUUUGG | |
| mecciND5-siRNA-2 | CUCCAACAUACUCGGAUUCU | AGAAUCCGAGUAUGUUGGAG | |
| mecciND5-siRNA-1 | CUACUCCCCCUAGACCUCAA | UUGAGGUCUAGGGGGAGUAG | |
| si-PNPASE | GCAGGUAGAAUUCCCACAA | UUGUGGGAAUUCUACCUGC | |
| mecciND1-AMO | TATGAGATTGTTTGGGCTACTGCTC | | |
| mecciND5-AMO | TTTTGGCTCGTAAGAAGGCCTAGAT | | |
| **qPCR primers for mecciRNA in 293T cells** | | | |
| 293Tmecci1_F | CAGCGCAATCCTATTCTAGA | | We chose top 50 mecciRNAs from 293T mitochondrial RNA-seq data, and 28 mecciRNA primers are suitable for qPCR detection. These primers are used for checking mecciRNA enrichment in FLAG-RIP experiments (Fig. 5F) |
| 293Tmecci1_R | GTAACTTGTTCCGTTGGTCA | | |
| 293Tmecci2_F | ACCTGGCGCAATAGATATAG | | |
| 293Tmecci2_R | GGGTAAATGGTTTGGCTAAG | | |
| 293Tmecci3_F | GCTGGTTGTCCAAGATAG | | |
| 293Tmecci3_R | TTGCTACATAGACGGGTG | | |
| 293Tmecci5_F1 | CAATCCTACCTCCATCGCTA | | |
| 293Tmecci5_R1 | AGGAGTAGGGTTAGGATGAG | | |
| 293Tmecci6_F | CTGTTAGTCCAAAGAGGAACAGC | | |
| 293Tmecci6_R | CAAGGGGATTTAGAGGGTTCTG | | |
| 293Tmecci7_F | TCACAGCACCAAATCTCCAC | | |
| 293Tmecci7_R | TTGTGCGGTGTGTGATGCTA | | |
| 293Tmecci8_F1 | CACACCCGTCTATGTAGCAA | | |
| 293Tmecci8_R1 | AGCTGTTCTTAGGTAGCTCG | | |
| 293Tmecci10_F1 | TACTCTTTCACCCACAGCA | | |
| 293Tmecci10_R1 | GAATCCGAGTATGTTGGAG | | |
| 293Tmecci15_F1 | TAGCATCACACACCGCACAA | | |
| 293Tmecci15_R1 | AGAATCCGAGTATGTTGGAG | | |
| 293Tmecci17_F1 | GACGTTAGGTCAAGGTGTAG | | |
| 293Tmecci17_R1 | AAGAGGTGGTGAGGTTGATC | | |
| 293Tmecci19_F | CACTCATCCTAACCCTACTC | | |
| 293Tmecci19_R | TTGGGTTGAGGTGATGATGG | | |
| 293Tmecci20_F | CCTAGACCTAACCTGACTAG | | |
| 293Tmecci20_R | GCTCGTAAGAAGGCCTAGAT | | |
| 293Tmecci22_F | CTCACTGTCAACCCAACACA | | |
| 293Tmecci22_R | TCTTAGGTAGCTCGTCTGGT | | |
| 293Tmecci23_F | CCAGACAACCTTAGCCAAAC | | |
| 293Tmecci23_R | CAAGAGGTGGTGAGGTTGAT | | |
| 293Tmecci24_F | CATGAGGTGGCAAGAAATGG | | |
| 293Tmecci24_R | CTTGCGCTTACTTTGTAGCC | | |
| 293Tmecci25_F | ATTCCGCTACGACCAACTCA | | |
| 293Tmecci25_R | AGAAGTAGGGTCTTGGTGAC | | |
| 293Tmecci26_F | ATTCTCCTCCGCATAAGC | | |
| 293Tmecci26_R | AGGGTGATAGATTGGTCC | | |
| 293Tmecci28_F | CACCAAATCTCCACCTCCATCAT | | |
| 293Tmecci28_R | CGGTGTGTGATGCTAGGGTA | | |
| 293Tmecci29_F | ATCTCGAACTGACACTGAGC | | |
| 293Tmecci29_R | CTTCTTGGTCTAGGCACATG | | |
| 293Tmecci30_F | CACTGTCAACCCAACACAG | | |
| 293Tmecci30_R | TCCTAGTGTCCAAAGAGCTG | | |
| 293Tmecci31_F | TCACAGCACCAAATCTCCAC | | |
| 293Tmecci31_R | CTAGTCAGGTTAGGTCTAGG | | |
| 293Tmecci32_F | CCGTACATAGCACATTACAG | | |
| 293Tmecci32_R | TGTACTACAGGTGGTCAAGT | | |
| 293Tmecci33_F | GGTCCATCATCCACAACCTT | | |
| 293Tmecci33_R | GTATGGCTTTGAAGAAGGCG | | |
| 293Tmecci34_F | CTCCTTACACTATTCCTCATCAC | | |
| 293Tmecci34_R | GGCATTTCACTGTAAAGAGGTGT | | |
| 293Tmecci35_F | ACCTTAGCTCTCACCATC | | |
| 293Tmecci35_R | CGTTCGGTAAGCATTAGG | | |
| 293Tmecci36_F | TCACACGATTAACCCAAGTCA | | |
| 293Tmecci36_R | CTGGCACGAAATTGACCAAC | | |

| | |
|---|---|
| 293Tmecci38_F | CCACAGGTCCTAAACTACCA |
| 293Tmecci38_R | AGAGCTGTTCCTCTTTGGAC |
| 293Tmecci39_F | CCTATCTAGGCCTTCTTACG |
| 293Tmecci39_R | TCTAGGGCTGTTAGAAGTCC |
